# Automatic inference of demographic parameters using Generative Adversarial Networks

**DOI:** 10.1101/2020.08.05.237834

**Authors:** Zhanpeng Wang, Jiaping Wang, Michael Kourakos, Nhung Hoang, Hyong Hark Lee, Iain Mathieson, Sara Mathieson

**Affiliations:** Department of Computer Science, Haverford College, Haverford, PA; Department of Computer Science, Swarthmore College, Swarthmore, PA; Department of Genetics, University of Pennsylvania, Philadelphia, PA

**Author notes:** Corresponding author: Sara Mathieson.

**Keywords:** Evolutionary modeling, Demographic inference, Generative adversarial network, Simulated data

## Abstract

Population genetics relies heavily on simulated data for validation, inference, and intuition. In particular, since the evolutionary “ground truth” for real data is always limited, simulated data is crucial for training supervised machine learning methods. Simulation software can accurately model evolutionary processes, but requires many hand-selected input parameters. As a result, simulated data often fails to mirror the properties of real genetic data, which limits the scope of methods that rely on it. Here, we develop a novel approach to estimating parameters in population genetic models that automatically adapts to data from any population. Our method, `pg-gan`, is based on a generative adversarial network that gradually learns to generate realistic synthetic data. We demonstrate that our method is able to recover input parameters in a simulated isolation-with-migration model. We then apply our method to human data from the 1000 Genomes Project, and show that we can accurately recapitulate the features of real data.

## Introduction

Simulation is a key component of population genetics. It helps to train our intuition, and is important for the development, testing, and comparison of inference methods. Because population genetic models such as the ancestral recombination and selection graphs [1, 2] are computationally intractable for inference but relatively easy to simulate, simulations are also heavily used for parameter inference. Approximate Bayesian Computation (ABC) [3] is a widely used example. Regardless of the application, the goal is to simulate data that is “realistic” in the sense that it resembles real data from the population(s) of interest. Typically this is done by fixing some parameters that are fairly well-known, then choosing other parameters to match some property of the real data, usually based on summary statistics. However, this involves a potential loss of information in the reduction to summary statistics and then an implicit weighting on the relative importance of different summary statistics. Often, parameters that create simulations that match one type of summary statistic (e.g. the site frequency spectrum) do not match others (e.g. linkage disequilibrium patterns) [4]. Here, we present a novel parameter learning approach using Generative Adversarial Networks (GANs). Our approach creates both realistic simulated data and a quantitative way of determining the match between any simulations proposed for a particular real dataset. For us, “realistic” means “cannot be distinguished from real data by a machine learning algorithm”, specifically a convolutional neural network (CNN).

Machine learning (ML) methods have been emerging more broadly as promising frameworks for population genetic inference. The high-level goal of training a ML method is to learn a function from the input (genetic data) to the output (evolutionary parameters). Some early efforts used machine learning to account for issues that arise with high-dimensional summary statistics [5–7]. More recently, machine learning approaches have used various forms of convolutional, recurrent, and “deep” neural networks to improve inference and visualization [8–14]. One of the goals of moving to these approaches was to enable inference frameworks to operate on the “raw” data (genotype matrices), which avoids the loss of information that comes from reducing genotypes to summary statistics. However, these algorithms rely heavily on simulated datasets for training. In machine learning more broadly, data is often hand-labeled with “true” values – part of this data is used to train the model, and part is held aside to test the model. In population genetics, such “labeled” training data is extremely limited because the evolutionary ground truth is rarely known with certainty. Thus all approaches rely on simulations to train and validate ML models.

Current simulators [15–21] are well equipped to replicate mechanisms of evolution, but require many user-selected input parameters including mutation rates, recombination and gene conversion rates, population size changes, natural selection, migration rates, and admixture proportions. We do not always have a good sense of what these parameters should be, especially in understudied populations and non-model species. For example, mutation and recombination rates estimated in one population are frequently used to simulate data for another, despite the fact that these rates differ between populations [22–26].

Generative models provide one route to simulating more realistic population genetic data. Typically, generative models create artificial data based directly on observed data, without an explicit underlying model. They have been used to create synthetic examples in a wide range of fields, from images and natural language to mutational effects [27] and single cell sequencing [28]. In particular, Generative Adversarial Networks (GANs) work by creating two networks that are trained together [29, 30]. One network (the *generator*) generates simulated data, while the other network (the *discriminator*) attempts to distinguish between “real” data and “fake” (synthetic) simulations. As training progresses, the generator learns more about the real data and gets better at creating realistic examples, while the discriminator learns to pick up on subtle differences and gets better at distinguishing examples. After training is complete, the generator can be used to create new examples that are indistinguishable (by the discriminator) from real data, but where the ground truth is known (i.e. labeled data).

The use of GANs in population genetics is just beginning. Recently Yelmen et al. [31] created a GAN that generates artificial genomes that mirror the properties of real genomes. Their approach does not include an evolutionary model, so the resulting artificial genomes are “unlabeled”. Such an approach is useful for creating proxy genomes that preserve privacy but still maintain realistic aggregate properties. However this synthetic data could not be used downstream to train or validate supervised machine learning methods since no evolutionary ground truth is known.

Here we present a parametric GAN framework that combines the ability to create realistic data with the interpretability that comes from an explicit model of evolution. The discriminator is a permutation-invariant CNN that takes as input a genotype matrix (representing a genomic region) and classifies it as real data or synthetic data. Throughout training, the discriminator tries to get better at this binary classification task. The generator is a coalescent simulator that generates genotype data from a parameterized demographic history. The generator is trained using a simulated annealing algorithm that proposes parameter updates leading to more discriminator confusion. The discriminator is trained using a gradient descent approach that is standard for neural networks. We apply our method, called `pg-gan`, in a variety of scenarios to demonstrate that it is able to recapitulate the features of real genetic data and confuse a trained discriminator. Although we focus on humans, the underlying methodology enables the simulation of any population or species, regardless of how much is known *a priori* about their specific evolutionary parameters.

We anticipate that the approach outlined in this work will be useful in strengthening the match between simulated and real data, especially for understudied populations that deviate from broad geographic groups. In addition, our discriminator can be used on its own (after training) to evaluate and compare different candidate simulations for the same real dataset. Downstream, our simulations can be used as a starting point for other methods that seek to quantify local evolutionary forces such as natural selection or mutation rate heterogeneity. There has also been a push in the population genetics community to standardize simulation resources [32] – we see our method as contributing to the assessment and refinement of published models as they are applied to new datasets.

## Materials and Methods

At a high level, our method works by simulating data from an underlying evolutionary model, then comparing it to real data via a neural network discriminator. As the discriminator is trained, it tries to minimize a loss function that incentivizes learning the difference between real data and synthetic data. But at the same time, the generator refines the evolutionary model so that it recapitulates the features of real data and attempts confuse the discriminator. At the end, the evolutionary model can be used to simulate additional realistic data for use in downstream applications or method comparisons. Additionally, the final parameters of the evolutionary model can be interpreted to learn more about the population or species of interest.

A GAN is not a traditional optimization problem – due to the dual nature of the generator and discriminator there are two optimization problems in a minimax framework, and it is difficult to evaluate the final trained model. Often the “GAN confusion” (discriminator classification accuracy) can be used to assess the success of the algorithm – a high classification accuracy (close to 1) indicates that the simulations are not capturing the real data and the discriminator is easily able to tell the difference between the two types of data. A low classification accuracy (close to 0.5) ideally indicates the evolutionary model has created simulations that are well-matched to the real data. However, an accuracy close to 0.5 could also mean that the discriminator has not learned anything and is either flipping a coin when classifying examples, or classifying all examples as the same class.

Training a GAN is a delicate balance. If the discriminator learns too quickly and becomes very good at identifying a specific setting of the simulated data from the real, then all proposals by the generator may look equally confusing. As a result, many generator proposals will be rejected and the discriminator will simply keep getting better a distinguishing the current setting from real data. On the other hand, if the discriminator learns too slowly, it may not be able to identify any generator proposals as better or worse. This often leads to a “random walk” across the parameter space, with the discriminator classifying everything as real or everything as simulated regardless of the generator’s proposals. Throughout the Methods section we outline techniques and strategies for balancing training and identifying degenerate states.

In the Method subsections below we first outline the notation for `pg-gan` and discuss the general training strategy. Then we provide further details about the generator and discriminator architectures. Finally we discuss applications of `pg-gan` to both simulated and real training data, as well as methods for evaluating the performance.

There are two inputs to the method, shown in orange in Figure 1. The first input is an evolutionary model parameterized by vector Θ. The parameters can be very flexible, including evolutionary event times, effective population sizes, and rates of mutation, recombination, migration, and exponential growth. The parameters Θ can be fed into the generator *G* to produce a simulated region *z*, which we write as

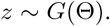

**Figure 1:**
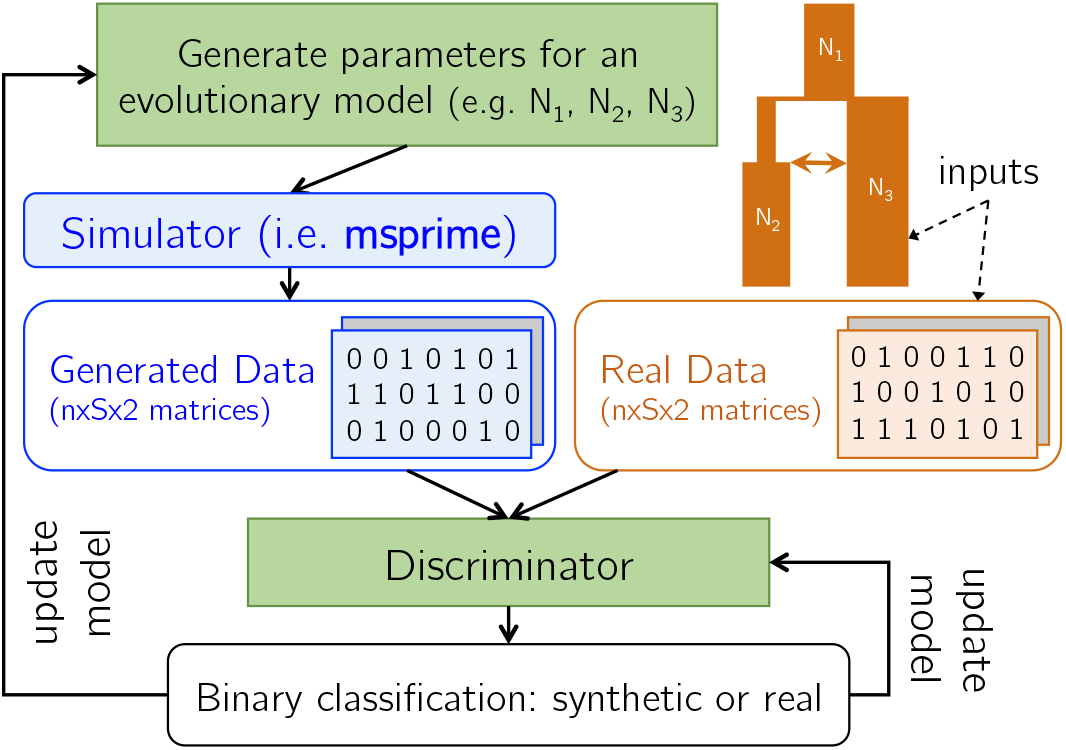
pg-gan algorithm overview. The inputs to our method are an evolutionary model and a set of real data (orange). The parameters of the generator and discriminator (green) are updated in a unified training framework using simulated annealing (generator) and backpropagation (discriminator). The generated data and real data are analyzed one genotype matrix at a time, where *n* is the number of haplotypes and *S* is the number of SNPs retained in each region. Inter-SNP distances are also fed in as a second channel, which provides the discriminator with information about SNP density.

The second input is a set of real data. We use *x* to denote a generic region from the real data. Both *z* and *x* have the same shape (*n, S,* 2) where *n* is the number of haplotypes, *S* is the number of SNPs in the region. The first channel represents the genotypes and the second channel represents the inter-SNP distances. The outputs of `pg-gan` are the optimal evolutionary parameters Θ* for the generator *G*, and a binary classifier *D* (the discriminator) which can predict if genomic regions are real or fake. Specifically, *D*(*x*) is the predicted probability that region *x* is *real*.

To incentivize the competing goals of the generator and discriminator, we minimize binary cross-entropy loss functions. If we have *M* regions of simulated data *{z*^(1)^*, …, z*^(*M*)^*}* generated under *G*(Θ), then the generator loss function is

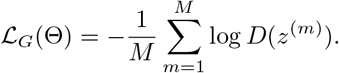

This loss function is cross-entropy, but where we only have one class (the generated data), which we want the discriminator to classify as real (label 1).

At the same time, the discriminator *D* is trying to classify the generated regions as fake (label 0) and the real regions as label 1. Therefore the discriminator loss function for *M* regions of real data *X* = *{x*^(1)^*, …, x*^(*M*)^*}* and *M* regions of simulated data *{z*^(1)^*, …, z*^(*M*)^*}* generated under *G*(Θ) is

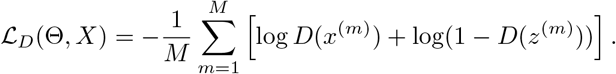

Algorithm 1 (in the style of [29]) shows the overall training of `pg-gan`.

**Algorithm 1:**
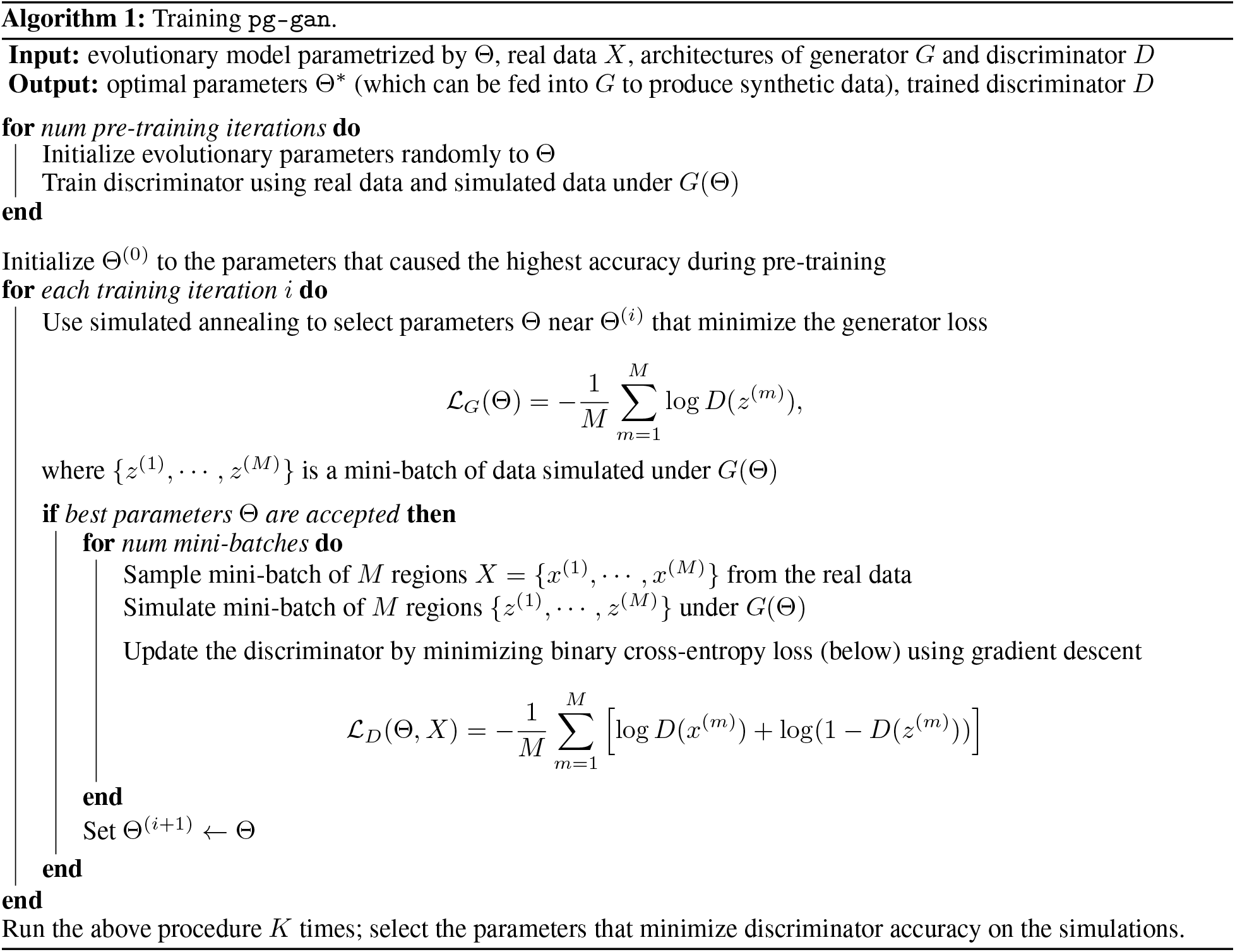
Training `pg-gan`.

### Generator

In image and video generation, the generator often takes the form of a CNN, since a large array of pixel information must be generated from a low-dimensional vector of noise (see Figure 1 of [33] for the architecture of a CNN-based image generator). For our purposes, we do not need to generate the individual genotypes for each training example, but we do need to generate candidate parameters for input into an evolutionary simulator (we use `msprime` [20] in this study).

Using this lens, we can view the generator learning problem as minimizing the multivariate generator loss function 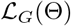 with respect to Θ. We optimize the loss using simulated annealing [34] due to its flexible parameter updates and lack of reliance on an analytic gradient. In simulated annealing, initial parameter values are proposed and then gradually refined. A *temperature* is used to control whether or not new parameter proposals are accepted. The temperature usually begins at a high value, indicating that sub-optimal parameter choices may be accepted liberally to facilitate exploration of the parameter space. As training proceeds, the temperature “cools”, reducing the chance of accepting a poor parameter choice and allowing the method to converge on a set of parameters that optimizes the desired function. Unlike ABC methods which require simulating from the entire parameter space before analyzing the real data, this simulated annealing approach uses the real data to adaptively narrow the focus to promising regions of the search space.

We use a pre-training phase (described in the **Discriminator** subsection) to choose a starting value for each evolutionary parameter, which forms the initial parameter vector Θ^(0)^. We set the temperature for simulated annealing *T* ^(0)^ = 1 and linearly decrease it to 0 over a fixed number of iterations. During each training iteration *i*, several new sets of candidate parameters are proposed, and evaluated based on the generator loss function 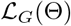. Each new set of parameters is proposed by sampling from a normal distribution around each current value, with variance based on the temperature. This allows the algorithm to explore the parameter space quickly in the beginning, and refine the estimates toward the end of GAN training. More formally, at iteration *i*, the candidate proposal for parameter *p* is

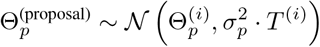

where 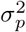 is the initial variance, which is based on the range of plausible values for each parameter. Out of the several candidate proposals, we choose the one that minimizes 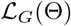. Then we compare this loss to the loss of the previous iteration. If the proposal reduces or maintains the generator loss, we always accept it. If not, we use the simulated annealing temperature to help define a threshold for acceptance. Formally, if the proposal is Θ and the current set of parameter values at iteration *i* is Θ^(*i*)^, then the acceptance probability is

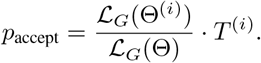

If the proposed parameters are accepted we train the discriminator using several mini-batches (with the simulated regions generated under Θ), then we set Θ^(*i*+1)^ *←* Θ. An important point is that we do not train the discriminator using the new parameter proposal unless it is accepted. During the candidate proposal phase, we are evaluating the parameter choices through the generator loss only.

### Discriminator

For the architecture of the discriminator, we use a permutation-invariant CNN based on `defiNETti` [8]. Each region *x* (real) or *z* (simulated) has shape (*n, S,* 2) where *n* is the number of haplotypes in the sample, *S* is the number of retained SNPs, and 2 indicates there is one channel for the genotypes and one channel for inter-SNP distances. The inter-SNP distances are duplicated down each column to allow this slice of the tensor to have the same shape as the genotype information. This also ensures that each convolutional filter processes the genotypes and associated distances at the same time. Alternatively, the convolutional layers can be used on the genotypes only, and the distances concatenated later as a vector. However, this approach does not allow the processing of the two channels to be as tightly coupled. We use convolutional filters of shape 1 *×* 5 (1 haplotype, 5 SNPs) to ensure that the order of haplotypes does not impact the results. We use ReLU as the activation function for all layers, and also use dropout [35] during training to guard against overfitting. After several convolutional layers, we condense the output by applying a column-wise permutation-invariant function. We experimented with both `max` and `sum` as permutation-invariant functions, and decided to use `sum` throughout. It generally causes the discriminator to learn more slowly than `max`, allowing the generator time to find good parameter choices. `max` sometimes causes the discriminator to converge quickly, easily distinguishing the real from simulated data before the generator can move to a promising location in the parameter space. Note that we need a fixed number of SNPs in each region to make sure the discriminator output is always of the same size. However, we do not need a consistent number of haplotypes, provided that the permutation-invariant function used is not sensitive to this number (i.e. `max` or `avg` would be fine but `sum` would not).

For models that consider multiple populations, we augment this framework to include separate permutation-invariant components for each population, then concatenate the flattened output before input into the dense layers at the end of the network. An illustration of our discriminator architecture for two populations is shown in Figure 2.

**Figure 2:**
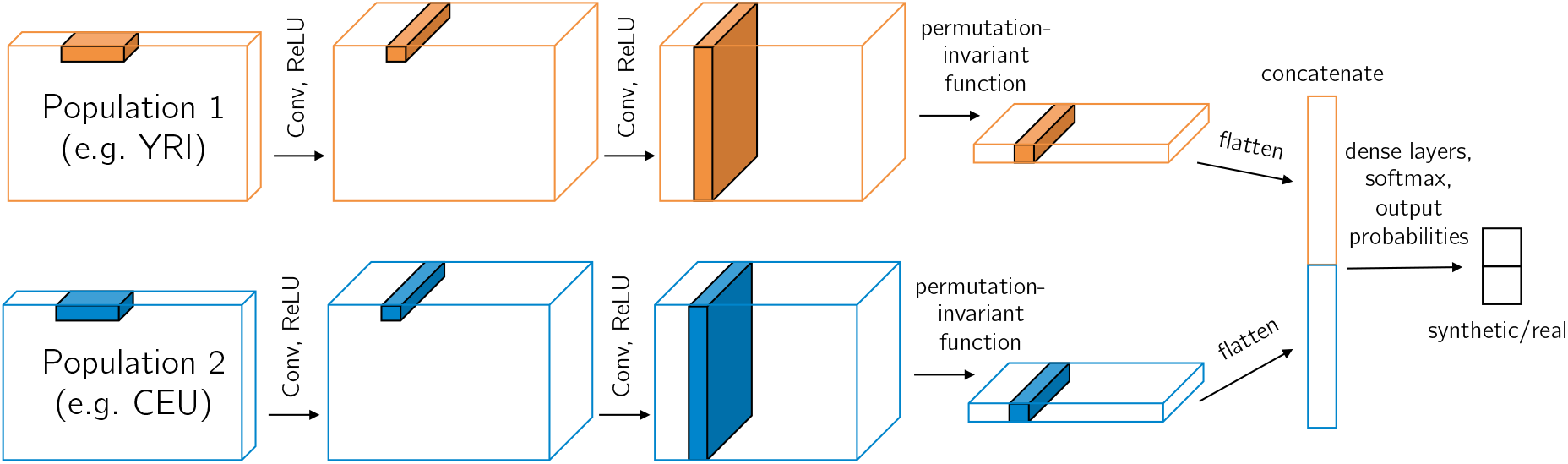
Multi-population CNN discriminator architecture. Each example region is of shape (*n, S,* 2) where *n* is the number of haplotypes (usually with *n/*2 from population 1 and *n/*2 from population 2). The convolutional filters for population 1 and 2 are shared (i.e. not separate weights) so that haplotype commonalities can be more easily identified. The final output of the discriminator is the probability the region is real (which can be subtracted from 1 to find the probability the region is simulated). This CNN can be reduced for one population or extended for three populations.

Through discriminator training we seek to minimize the loss function 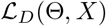, with a small entropy term subtracted to disincentivize predicting all the same class. This entropy term is different from the entropy regularization used to prevent mode collapse [36], a common problem in GAN training. In such cases, the goal is to increase the entropy of the *generator* so it can produce a multi-modal distribution (e.g. different types of images such as hand-written digits). Mode collapse is not an issue for `pg-gan`, as a single set of evolutionary parameters is desired. To minimize our discriminator loss function we use gradient descent (via backpropagation) with mini-batches of 100 training examples (half are real and half are simulated). For each training iteration, we perform 100 mini-batch training updates if the proposed parameters are accepted. This allows the discriminator to learn gradually, as the parameters are being refined. While a classification accuracy close to 0.5 is desired by the end of training, the discriminator accuracy may be close to this value early on in the training process simply because it has not learned anything yet. The goal is for the discriminator to be optimized to distinguish real from simulated data as much as possible and *still* be wrong half the time.

Due to the simulated annealing training of the generator, initial step sizes of the parameters can be large to explore the parameter space more quickly. This can present a problem for discriminator training. If the parameters change too quickly, the discriminator does not have time to learn the difference between the real data and data simulated under a wide variety of parameter setting. In some situations, this leads the discriminator to fail to learn anything and it predicts the same class (either real or fake) for all regions. To combat this issue, before GAN training begins we pre-train the discriminator only, using a variety of randomly sampled parameter values. We find that pre-training gives the discriminator an overall sense of the data, so that when generator training begins the discriminator is able to identify which generated regions were closer to the real and which were further away. We run the combined (generator and discriminator) training for 300 iterations.

### Simulation study

To validate our approach, we first select the training dataset to be a simulated one, so that we can test whether the inferred parameters are correct. To assess a variety of different types of parameters, we choose an isolation-with-migration model (see Figure 3A) with six parameters. The parameters include three effective population sizes: *N*_anc_ for the ancestral population size, and *N*_1_ and *N*_2_ for the sizes of each population after the split. We also infer the split time *T*_split_, and the strength of a directional migration pulse (mig) at time *T*_split_*/*2. Finally, we infer the per-base, per-generation recombination rate (reco). We evaluate the inferred parameters based on how well they match the true parameters. See Table 1 for the ranges and units of each parameter.

**Table 1:** Parameter ranges. When inferring a parameter, we initialize its value by drawing a value uniformly from the given ranges. For each parameter update, we do not allow the parameter to go up to or outside its range. Overall the ranges are meant to be plausible values based on previous studies or reasonable evolutionary events.

| Parameter | min | max | units |
| --- | --- | --- | --- |
| $N_e$ | 1000 | 30000 | individuals |
| reco | $1 \times 10^{-9}$ | $1 \times 10^{-7}$ | per base per generation |
| mut | $1 \times 10^{-9}$ | $1 \times 10^{-7}$ | per base per generation |
| $N_{\text{anc}}$ | 1000 | 25000 | individuals |
| $T_{\text{split}}$ | 500 | 20000 | generations |
| mig | -0.2 | 0.2 | fraction of individuals |
| $N_1$ | 1000 | 30000 | individuals |
| $N_2$ | 1000 | 30000 | individuals |
| growth | 0 | 0.05 | per generation |
| $N_3$ | 1000 | 30000 | individuals |
| $T_1$ | 1500 | 5000 | generations |
| $T_2$ | 100 | 1500 | generations |

**Figure 3:**
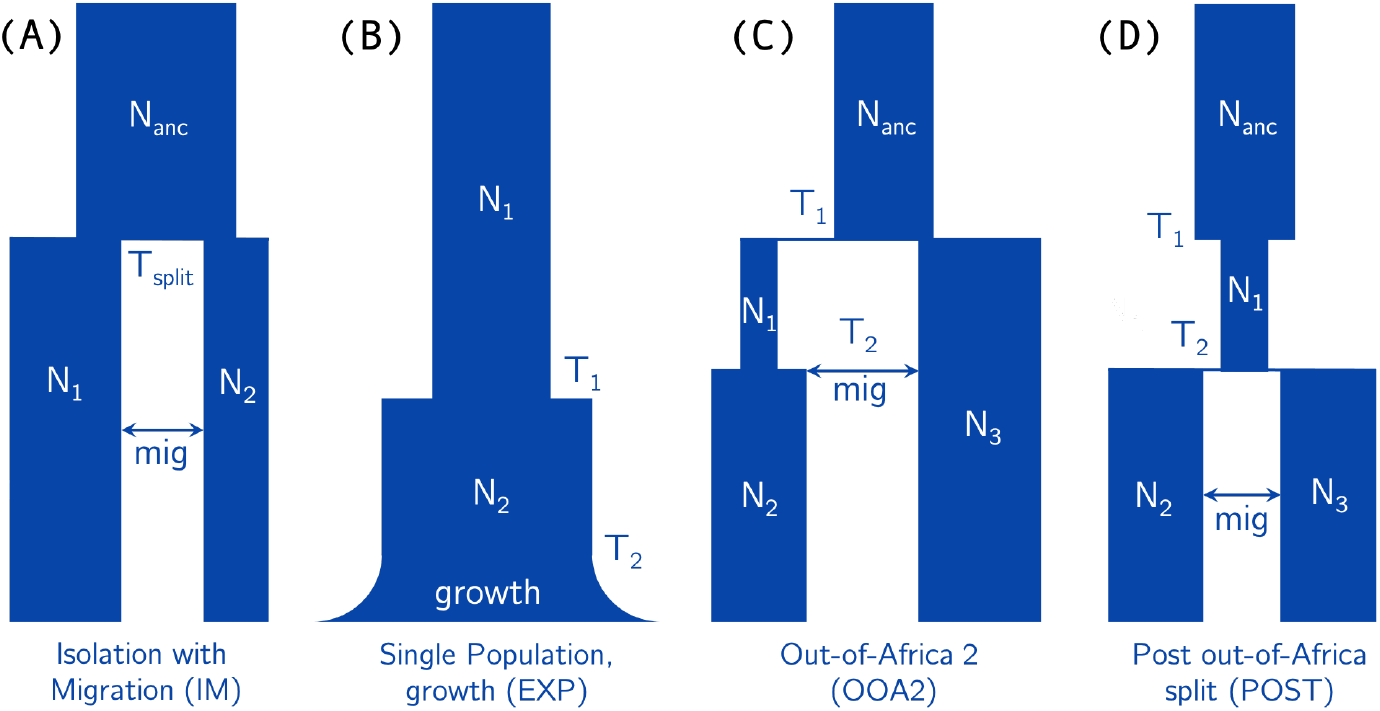
Set of models. (A) A six-parameter, two-population isolation-with-migration model, which we use in the simulation study. The migration event is a single pulse at time *T*_split_*/*2, and can be in either direction. The final parameter (not shown in this diagram) is the recombination rate. (B) A five-parameter, single-population exponential growth model, which we use to infer histories for YRI, CEU, and CHB separately. (C) A seven-parameter, two-population model, which we fit separately for YRI/CHB and YRI/CEU. The migration can be in either direction. (D) A seven-parameter, two population model which we fit to CEU/CHB. Migration occurs at *T*_2_*/*2 and can be in either direction.

### 1000 Genomes data analysis

To demonstrate the effectiveness of our method on real data, we use `pg-gan` to infer demographic parameters for both single- and multi-population models in humans. To ensure that the real data is as similar as possible to the simulated data, we run several pre-processing steps. To avoid processing the real data on-the-fly during training, we follow a data extraction pipeline to convert the real data into HDF5 format [37, 38]. Before converting VCF information into HDF5 format, we select haplotypes from each population and filter non-segregating and multi-allelic sites. The number of haplotypes is flexible (due to the permutation-invariant framework). We use between 196 and 198, matching the minimum number of individuals in each 1000 Genomes population.

During training, for each region *x* fed into `pg-gan`, we select a start SNP randomly from the entire genome. This random start point mitigates the effects of correlated nearby regions and local variations in mutation and recombination rate. Starting with this SNP, *S* = 36 biallelic SNPs are retained (along with their inter-SNP distances), which means the region has a flexible length. If 36 SNPs would cause the region to extend past the end of a chromosome, we reject the start SNP and sample a new one. For each region, we retain it if at least 50% of the bases are inside callable regions, as defined by the “20120824” strict mask [39].

For both the real and simulated data we recode the genotypes by setting the minor allele to the value “1” and the major allele to the value “*−*1” so that the discriminator cannot learn to distinguish real data based on reference bias or ancestrally misidentified states. For the simulations where we must specify a region length *L*, we choose *L* = 50kb, which ensures that in the majority of situations we have at least *S* = 36 SNPs. The middle 36 SNPs are retained, and any regions with insufficient SNPs are centered and zero-padded. Such regions would automatically look very different from the real data, so the generator quickly learns to avoid parameters that cause insufficient SNPs.

We test four models: Figure 3B-D, as well as a three-population Out-of-Africa model originally specified in [40]. The single-population model has five parameters: two effective population sizes *N*_1_ and *N*_2_, two size-change points *T*_1_ and *T*_2_, and the rate of exponential growth in the recent past. We fit this model to three human populations from the 1000 Genomes project: YRI (West African), CEU (European), and CHB (East Asian). The second model (OOA2) is a simplified two-population Out-of-Africa model. There are seven parameters: four effective population sizes, two time-change points, and a migration pulse that can be in either direction, allowing for migration between African and non-African populations. We fit this model to two pairs of populations: YRI/CEU and YRI/CHB. The third model (POST) represents the post-out-of-Africa split between the ancestors of Europeans and East Asians. In this seven-parameter model we allow a pre-split bottleneck and directional migration. We fit this model to the pair of populations CEU/CHB. Finally, we apply the three-population Out-of-Africa model (OOA3) to YRI/CEU/CHB, as implemented in `stdpopsim` [32].

### Evaluation metrics

One pervasive issue with GANs is the lack of a natural evaluation metric (see [41] for a comprehensive overview of GAN evaluation metrics). Many GANs have been evaluated qualitatively through user studies designed to see if humans find the generated data realistic [42]. For images, videos, or text this type of evaluation can be informative (although it tends to favor generators that memorize specific real examples [41]), but this is not directly possible in the case of genetic data.

Visualizing summary statistics is an alternative, although since we do not know which statistics are sufficient for the model, it is dangerous to rely on these alone as a final evaluation metric. It is possible the discriminator is learning other statistics or representations of the data that we are not aware of. In addition, explicitly matching some types of statistics can bias the resulting fitted model. For example, Beichman et al. [4] found that SFS-matching methods like *∂*a*∂*i [40] and `SMC`++ [43] are not able to recapitulate LD statistics. Further, we currently do not have an exhaustive or sufficient set of summary statistics that could be used to identify model parameters directly in a likelihood framework. However, as a qualitative assessment of our results, we compare summary statistics computed on the real data and data simulated under our inferred parameters. This gives us a sense of which features of real data agree with our simulations and which do not.

To that end, we use seven types of summary statistics. In all cases, we use 5000 regions of real data (chosen randomly) and 5000 regions of simulated data (each simulated independently under our inferred parameters) to compute the statistics. All pre-processing is the same as for GAN training, except for Tajima’s D where we fix the region length, not the number of SNPs.

- **SFS**: we compute the site frequency spectrum (SFS) by counting the number of singletons, doubletons, etc in each of 5000 regions of real and simulated data. We plot the first 10 entries.
- **Inter-SNP distances**: we plot the distribution of inter-SNP distances for both the real and simulated data (measured in base pairs). This provides a general measure of SNP density.
- **LD**: we compute linkage-disequilibrium (LD) by clustering pairs of SNPs based on their inter-SNP distance. We divide these distances into 15 bins and average the correlation *r*^2^ within each one.
- **Pairwise heterozygosity**: we plot the distribution of pairwise heterozygosity (*π*), computed separately for each region.
- **Tajima’s D**: we plot the distribution of Tajima’s D, computed separately for each region. Here we fix the region length to *L* = 50kb instead of fixing the number of SNPs, as otherwise the distribution would be the same as pairwise heterozygosity.
- **Number of haplotypes**: we plot the distribution of number of haplotypes for each region.
- **Hudson’s***F*_**st**_: for the two-population split models, we use *F*_st_ to measure population differentiation [44].

As a more quantitative evaluation, we also report the final discriminator classification accuracy. However, even this metric is not easy to interpret, as an accuracy close to 0.5 may indicate a degenerate situation where the discriminator has not learned anything (see Figure S1 for an example). Thus, for each model and dataset we run `pg-gan` *K* = 10 times and select the model that minimizes the classification accuracy of the discriminator on the final *generated* data (not using any real data). The more generated data that the discriminator classifies as real–i.e. the lower the discriminator accuracy–the better the generator. This metric was inspired by the Inception Score [45] used to evaluate GANs, where generated data is fed into a more powerful discriminator. Since no generated region has ever been seen by the discriminator before, all generated regions are implicitly “test data”. In this way we avoid relying on a held-aside real dataset for evaluation, which allows us to use the (limited) real data exclusively for training.

Finally, we also ran a comparison study against the ABC method `fastsimcoal` [46]. We provided `fastsimcoal` with three of the same models (IM, OOA2, and POST) as well as a simulated joint SFS, and the full genome joint SFS for YRI/CEU, YRI/CHB, and CEU/CHB. Then we compared the parameters and summary statistics from `fastsimcoal` to those from `pg-gan`.

Data Availability Statement: Our `pg-gan` software uses a `tensorflow` [47] backend and is available open-source at https://github.com/mathiesonlab/pg-gan. All data included in this work is publicly available through the 1000 Genomes Project https://www.internationalgenome.org/ [39].

## Results

### Simulation study

To validate our method, we first simulated the training dataset, so we knew the true evolutionary parameters. We fit the six-parameter IM model from Figure 3A, using the parameter ranges in Table 1. Throughout, we usually fix the mutation rate to 1.25 *×* 10^*−*8^ per base per generation, but it could be inferred along with the other parameters in species or populations where it is less established. See Figure S2 for an example where we infer mutation rate as well as the other six parameters of the IM model.

During the pre-training phase we train the discriminator on up to 10 different parameter sets, randomly chosen from the ranges in Table 1. We select Θ^(0)^ to be the first set that achieves at least 90% discriminator classification accuracy (or the set that maximizes accuracy in the case when we do not achieve 90% after 10 pre-training iterations). This enables the discriminator to gain some structure that is relevant to the data before combined training begins. During each main training iteration, we choose 10 independent proposals for each parameter, keeping the other parameters fixed. This creates 10 *× P* possible parameter sets, where *P* is the number of parameters (*P* = 6 for the IM model). We select the set that minimizes the generator loss, which has the effect of modifying one parameter each iteration. We also tested modifying all the parameters each iteration, but generally found that updating one at a time led to more stable and consistent results. For each parameter *p*, we set the initial variance 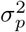 to the parameter range divided by 15.

We performed 10 independent initializations of `pg-gan` on the full set of six parameters for the IM model. We selected the results that minimized discriminator accuracy on the final generated data. The results are shown in Figures 4,5 and Table 2. The first subplot in Figure 4 shows the losses for both the generator and discriminator. Since the generator loss considers half as many regions, it is multiplied by two to be on same scale as the discriminator loss. At first the generator loss is high and the discriminator loss is low because the discriminator is easily able to detect the difference between simulated and training data. The second plot shows the discriminator accuracy on both simulated and training data. Both accuracies are initially high and then reduce to around 0.5. We see that here, `pg-gan` is able to find parameter values that bring the discriminator close to an accuracy of 0.5. The final classification accuracy on generated data here was 0.54, and the overall accuracy (considering both generated and training data) was 0.46. Our inferred parameter values are close to the true values (Table 2) and the site frequency spectrum and other summary statistics of data simulated with these parameter values closely match the summary statistics of the training data (Figure 5).

**Table 2:**
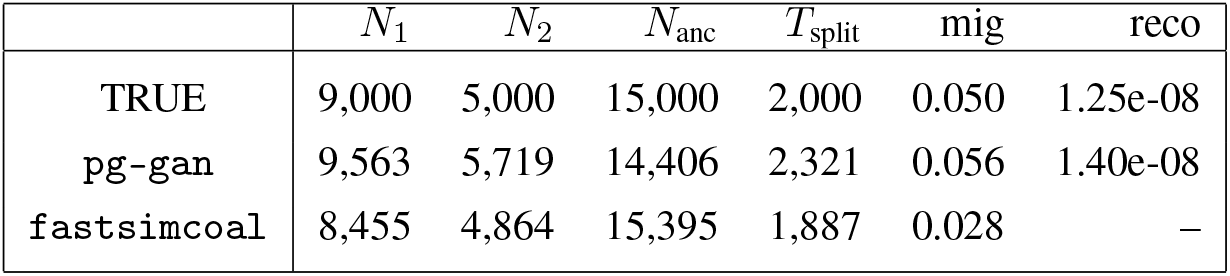
Comparison of pg-gan and fastsimcoal. Inferred parameters for the IM model (see Figure 3A). `pg-gan` results correspond to Figures 4,5, and `fastsimcoal` results correspond to Figure 6. For `fastsimcoal`, recombination rate is not shown, since it cannot be inferred from the SFS.

| | $N_1$ | $N_2$ | $N_{anc}$ | $T_{split}$ | mig | reco |
| --- | --- | --- | --- | --- | --- | --- |
| TRUE | 9,000 | 5,000 | 15,000 | 2,000 | 0.050 | 1.25e-08 |
| pg-gan | 9,563 | 5,719 | 14,406 | 2,321 | 0.056 | 1.40e-08 |
| fastsimcoal | 8,455 | 4,864 | 15,395 | 1,887 | 0.028 | — |

**Figure 4:**
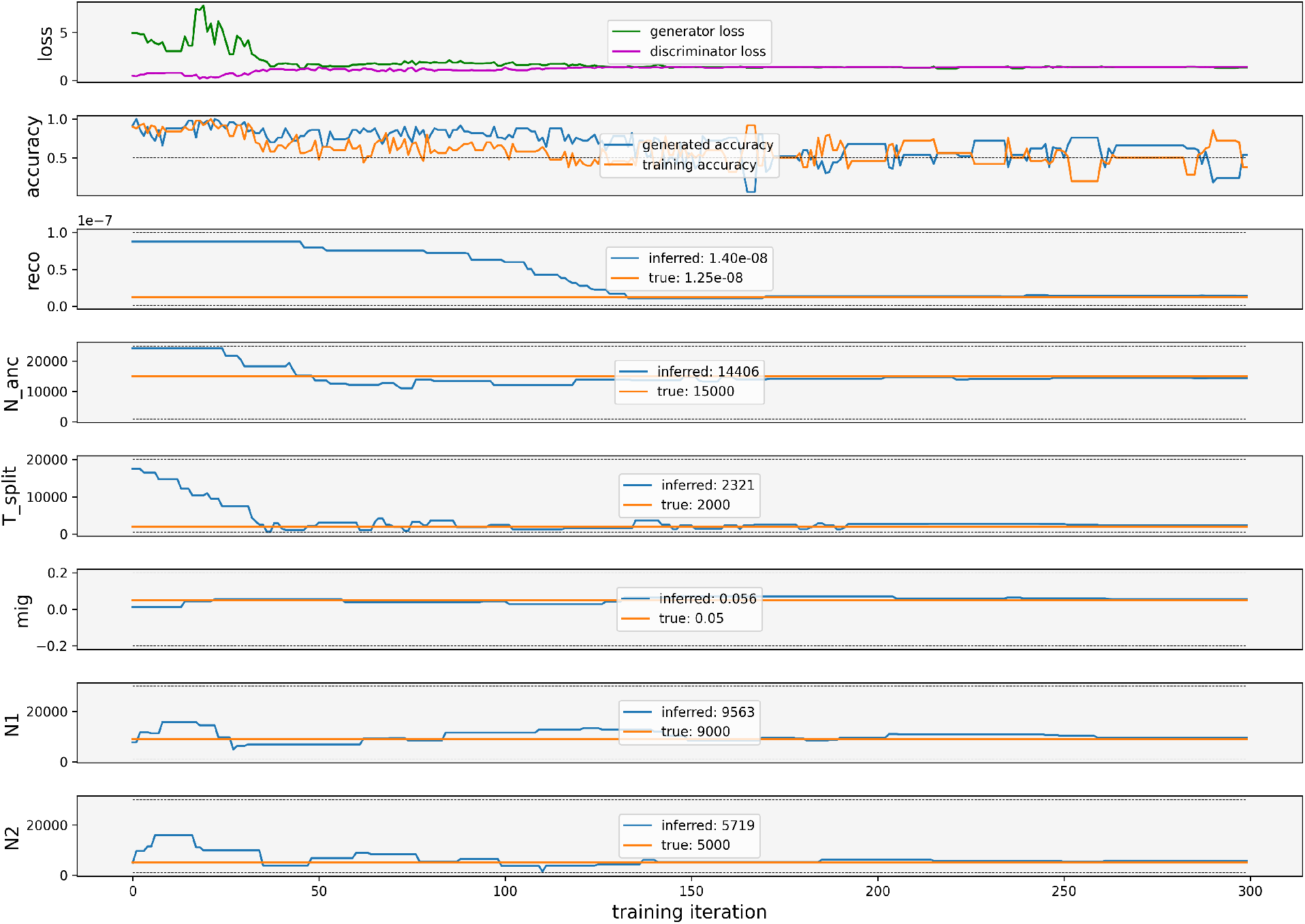
IM model parameter inference on simulated training data. In this scenario we jointly infer the six parameters of the IM model from Figure 3A. The top plot shows both loss functions over the course of GAN training, and the second plot shows classification accuracy for both simulated and training data. The remaining plots show the model parameters as they are refined throughout GAN training. The inferred values are taken at the final iteration.

**Figure 5:**
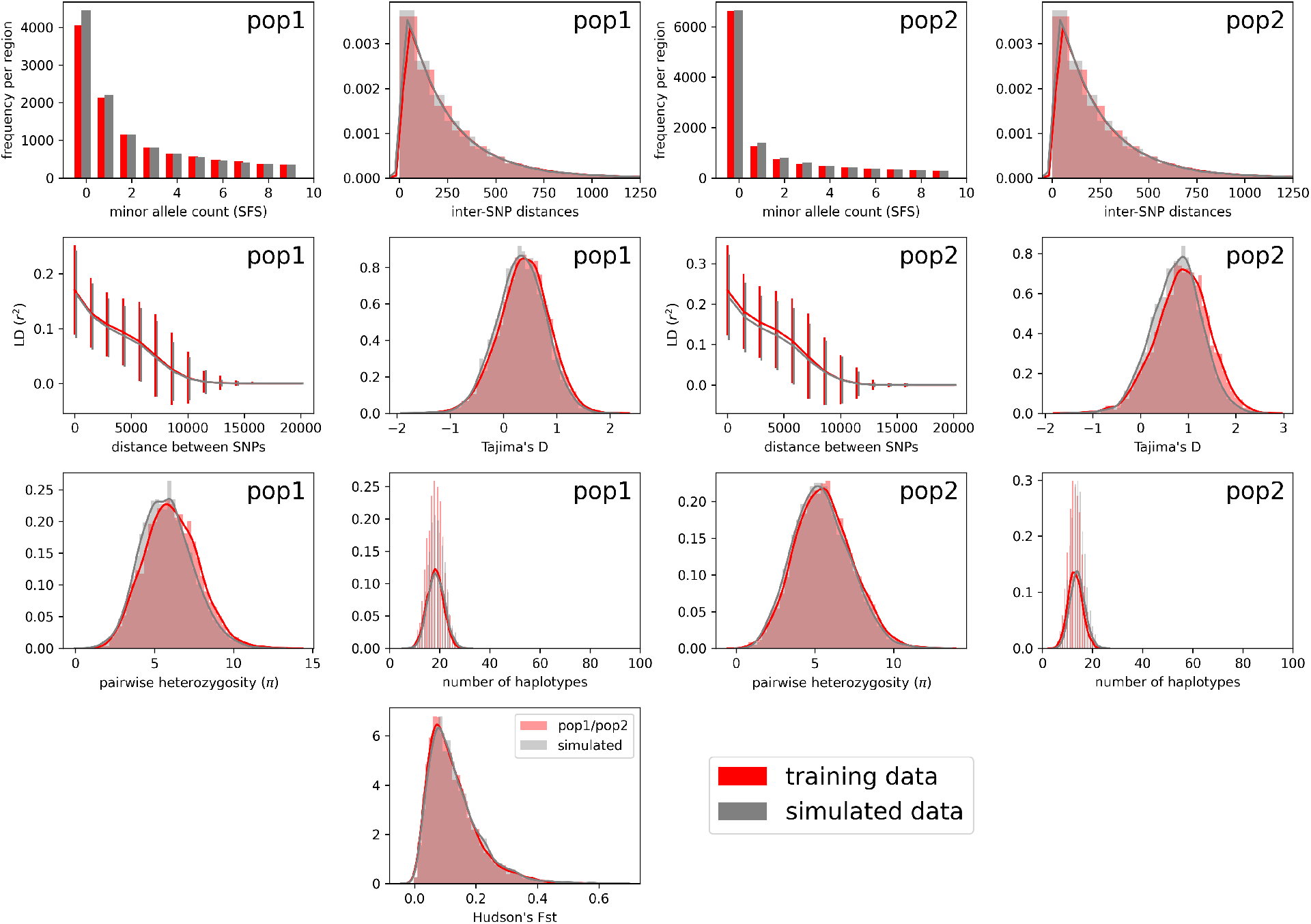
IM model statistics on simulated training data. Summary statistics for data simulated under our inferred parameters (“simulated data”), compared with data simulated under the true parameters (“training data”). Subfigures on the left correspond to statistics from the first population, and those on the right correspond to the second population. In the bottom panel we show *F*_st_ between the two populations.

As a comparison, we performed ABC inference using `fastsimcoal` using the same IM model, fitting the joint SFS from data simulated under the true model parameters. `fastsimcoal` closely matches the SFS and true parameters (Figure 6, Table 2), although it is not able to infer recombination rate. If we give it the correct recombination rate then it closely matches the other summary statistics in Figure 5 as well.

**Figure 6:**
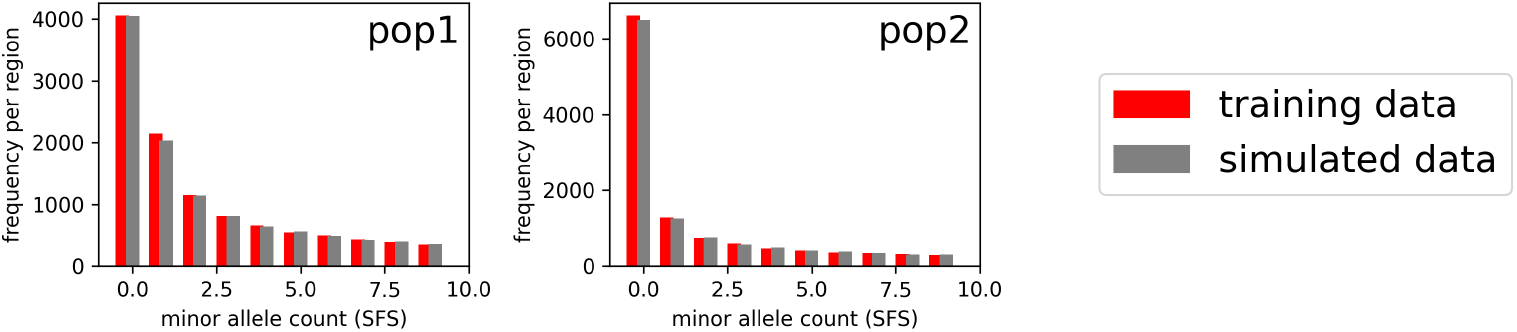
IM model SFS as inferred by fastsimcoal. Here we compare the true SFS (“training data”) with the SFS computed from data simulated under the parameters learned by `fastsimcoal` (“simulated data”).

### 1000 Genomes data analysis

We analyzed three populations (YRI, CEU, and CHB) separately, each under the five-parameter model with recent exponential growth (EXP; Figure 3B). For all single-population results we used *n* = 198 (size of CEU) and *S* = 36. Unlike the simulated example above, we fix the distribution of recombination rates by sampling from the distribution of HapMap combined recombination rates [48], although in principle this distribution could also be inferred.

To assess the impact of the model on our evaluation metrics (classification accuracy and summary statistics), we first fit a one-parameter demographic model with a single constant population size *N*_*e*_. We then contrast this result with the five-parameter exponential growth model (EXP). The summary statistics for these results are shown in Figure 7 for YRI and CHB, and a summary for all populations is shown in Figure 8A. Inferred parameters for each population under the five-parameter exponential growth model are shown in Table 3. The effect of the Out-of-Africa bottleneck (*N*_2_) is very apparent in CEU and CHB, but absent in YRI. Data simulated with fitted parameters for YRI contain many more singletons than the real data, possibly indicating the recent exponential growth rate (or the time of onset *T*_2_) is overestimated. On the other hand, low power to detect rare variants in the real data could explain a lack of singletons in YRI or other populations.

**Table 3:** 1000 Genomes single population parameter inference. Inferred parameters for the exponential growth model (see Figure 3B) in YRI, CEU, and CHB. We generally infer similar parameters for CEU and CHB.

| Population | $N_1$ | $N_2$ | growth | $T_1$ | $T_2$ |
| --- | --- | --- | --- | --- | --- |
| YRI | 23,676 | 20,837 | 0.0379 | 2,498 | 1,214 |
| CEU | 25,127 | 4,676 | 0.0061 | 1,673 | 949 |
| CHB | 21,136 | 3,150 | 0.0242 | 2,584 | 645 |

**Figure 7:**
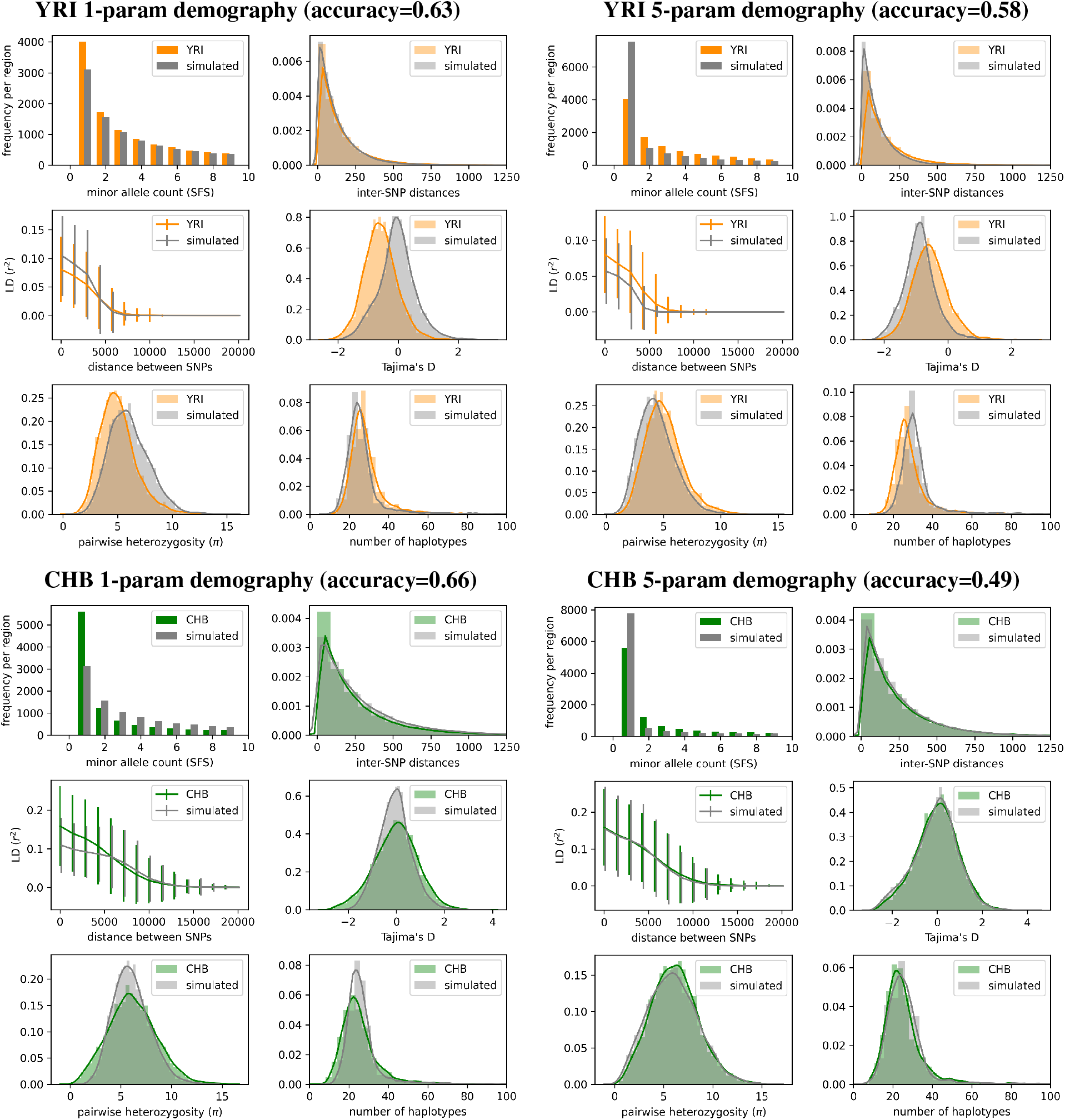
Single-population model. Summary statistic comparisons between 1000 Genomes Project data and data simulated under our `pg-gan` inferred parameters for a variety of scenarios. Top left: YRI vs. data simulated under the 1-parameter constant population size model. Simulated accuracy: 0.52, overall accuracy: 0.63. Top right: YRI vs. data simulated under the 5-parameter exponential growth model. Simulated accuracy: 0.72, overall accuracy: 0.58. Bottom left: CHB vs. data simulated under the 1-parameter constant population size model. Simulated accuracy: 0.68, overall accuracy: 0.66. Bottom right: CHB vs. data simulated under the 5-parameter exponential growth model. Simulated accuracy: 0.54, overall accuracy: 0.49.

**Figure 8:**
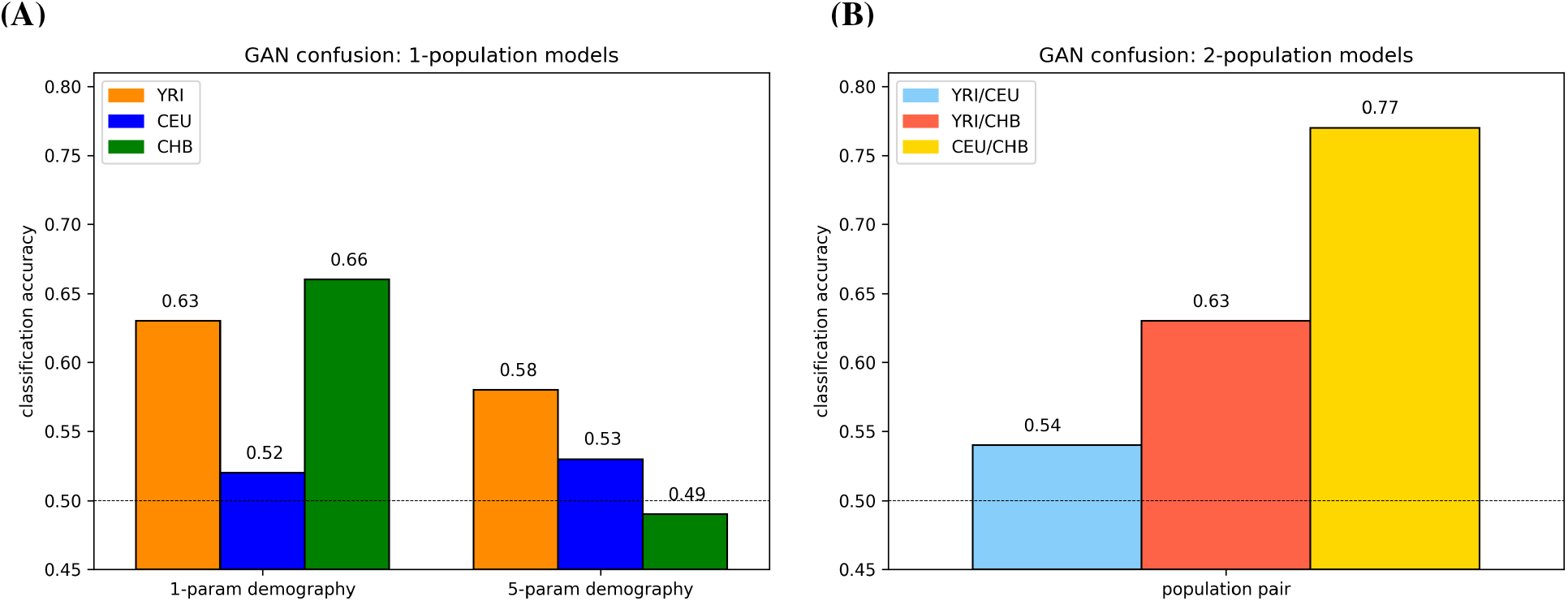
GAN confusion for 1- and 2-population models. (A) Comparison of 1- and 5-parameter models. We use a constant population size for the first group of bars, then move to the five-parameter exponential growth model (Figure 3B). We sample recombination rates from HapMap in both scenarios, instead of fixing the recombination rate. (B) Classification accuracy results on the population split models for YRI/CEU, YRI/CHB, and CEU/CHB. The Out-of-Africa models and parameter inference for YRI/CEU and YRI/CHB generally seem to do well, but the CEU/CHB split model and/or parameter inference does not result in simulated data that matches real data.

We also compared summary statistics (Methods) between the real data and data simulated under the parameter choices corresponding to the two scenarios from Figure 8A. In Figure 7 we show two sets of summary statistics each for YRI and CHB. On the left we show the one-parameter demography results, and on the right we show the 5-parameter results (using HapMap recombination rates in both cases). While some statistics match closely, others are less well-matched, consistent with the discriminator being imperfectly confused. Summary statistics for CEU are shown in Figure S3. For CEU both the 1- and 5-parameter models produced low classification accuracy, but the summary statistics are imperfect. This likely indicates that the discriminator did not learn as well in this scenario, not that the generator is producing high-quality simulated data.

For all our results, we discard any run where the discriminator classifies all regions in the same way (either all real or all simulated) at the end of training. For each set of 10 runs, 0-2 runs typically fail in this way. See Figure S1 for an example of a failed run for YRI. For the remaining runs, we see a range of final classifications accuracies. For the 5-parameter models, in YRI this range was 0.5–0.77 (mean 0.619) and for CHB this range was 0.49–0.67 (mean 0.564).

To pick the final result, we use the accuracy on the generated data only (i.e. not including the training data); 0.64–1.0 with mean 0.742 for YRI and 0.54-0.9 with mean 0.707 for CHB.

Next, we ran `pg-gan` on 1000 Genomes data from two populations. To model the split of African and non-African populations, we use two pairs of populations separately: YRI/CEU and YRI/CHB, using the OOA2 model from Figure 3C. We use CEU/CHB with the POST model from Figure 3D to represent the post-out-of-Africa split between the ancestors of Europeans and East Asians. The resulting classification accuracies are shown in Figure 8B. The YRI/CEU and YRI/CHB results are comparable to the single population analysis, but the CEU/CHB classification accuracy is much higher. For all pairs of populations, we provide the parameter inference results in Table 4. Summary statistics for the YRI/CEU split (Figure 9) match the real data closely. YRI/CHB statistics are shown in Figure S4 – for the YRI samples these statistics are not quite as closely matched, consistent with the slightly higher classification accuracy for this scenario. CEU/CHB statistics are shown in Figure S6 and are less well-matched to the real data, consistent with the relatively high classification accuracy, and suggesting that this model does not contain all the important features for these population, for example archaic admixture or exponential growth.

**Table 4:**
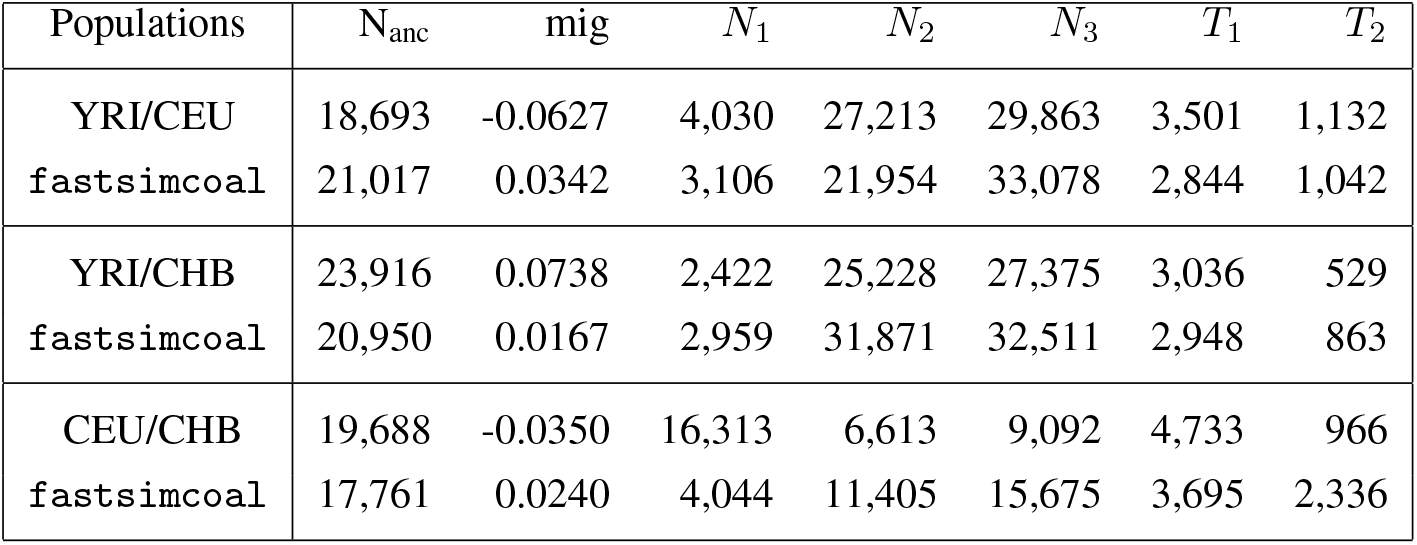
1000 Genomes two-population parameter inference. Inferred parameters for the OOA2 model (see Figure 3C) fit to YRI/CEU and YRI/CHB, as well as the POST model (see Figure 3D) fit to CEU/CHB. Results for both `pg-gan` and `fastsimcoal` are included. For `pg-gan`, we generally see similar results for YRI/CEU and YRI/CHB, with a lower classification accuracy for YRI/CEU, indicating a closer match to the real data. Our results are broadly consistent with `fastsimcoal`, except for CEU/CHB, where neither method produces statistics that match the real data (see Figures S6 and S7).

| Populations | $N_{\text{anc}}$ | mig | $N_1$ | $N_2$ | $N_3$ | $T_1$ | $T_2$ |
| --- | --- | --- | --- | --- | --- | --- | --- |
| YRI/CEU | 18,693 | -0.0627 | 4,030 | 27,213 | 29,863 | 3,501 | 1,132 |
| fastsimcoal | 21,017 | 0.0342 | 3,106 | 21,954 | 33,078 | 2,844 | 1,042 |
| YRI/CHB | 23,916 | 0.0738 | 2,422 | 25,228 | 27,375 | 3,036 | 529 |
| fastsimcoal | 20,950 | 0.0167 | 2,959 | 31,871 | 32,511 | 2,948 | 863 |
| CEU/CHB | 19,688 | -0.0350 | 16,313 | 6,613 | 9,092 | 4,733 | 966 |
| fastsimcoal | 17,761 | 0.0240 | 4,044 | 11,405 | 15,675 | 3,695 | 2,336 |

**Figure 9:**
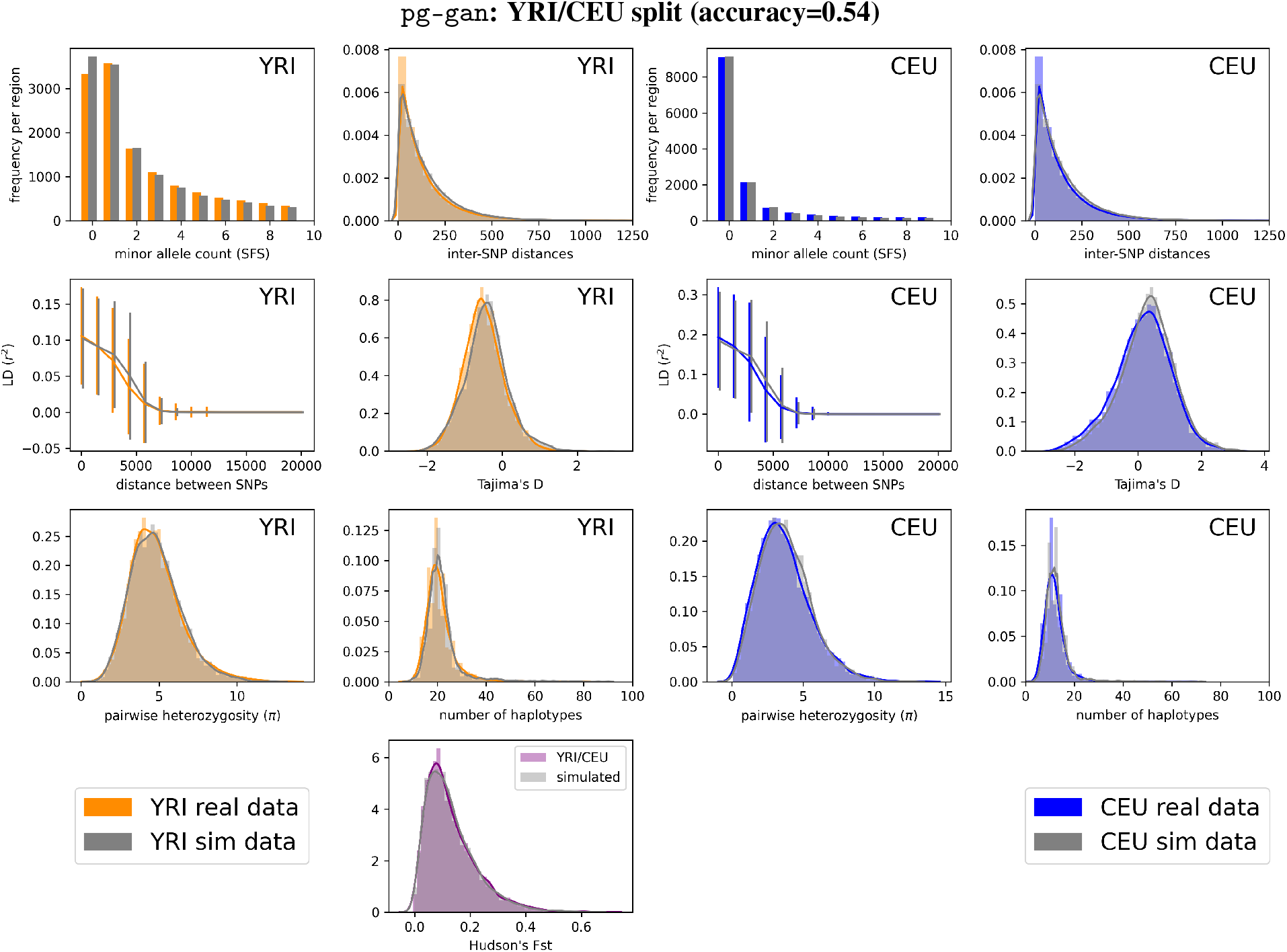
YRI/CEU: two-population model. Summary statistic comparison real 1000 Genomes data and data simulated under the inferred parameters from Table 4 (first row). Left: statistics computed on YRI samples only. Right: statistics computed on CEU samples only. Sites with count zero are segregating in only one population. *F*_st_ between the two populations is shown in the bottom panel. Simulated accuracy: 0.68, overall accuracy: 0.54.

We also ran `fastsimcoal` on the joint SFS from YRI/CEU, YRI/CHB (using the OOA2 model from Figure 3C), and CEU/CHB (using the POST model from Figure 3D). We used the inferred parameters to create new simulations (with the same fixed mutation rate and HapMap recombination rate distribution used for `pg-gan`). The resulting summary statistics for YRI/CEU are shown in Figure 10, demonstrating that `fastsimcoal` also matches the real data very well. The other `fastsimcoal` results are shown in Figure S5 (YRI/CHB) and Figure S7 (CEU/CHB). For YRI/CHB `fastsimcoal` produces a slightly better fit than `pg-gan`, but for CEU/CHB the two methods produce very different parameter estimates and neither method matches the summary statistics very well, supporting the suggestion that the generative model is missing some key features of the data.

**Figure 10:**
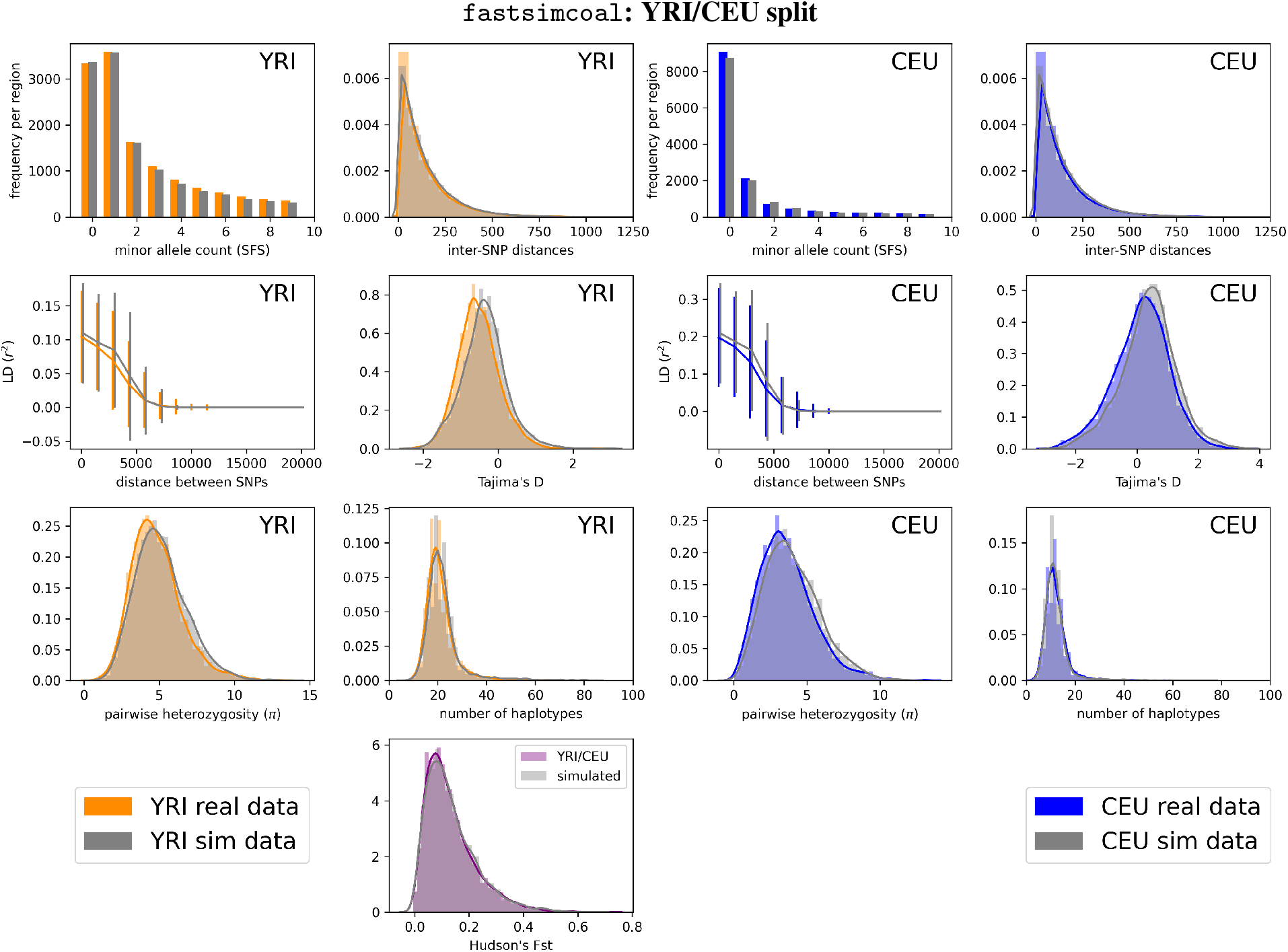
YRI/CEU: two-population model (fastsimcoal). Summary statistic comparison between YRI/CEU and data simulated under the OOA2 model parameters inferred by `fastsimcoal`. Here we include all the statistics (unlike Figure 6) since we are providing `fastsimcoal` with a recombination rate distribution. Left: statistics computed on YRI samples only. Right: statistics computed on CEU samples only. Sites with count zero are segregating in only one population. *F*_st_ between the two populations is shown in the bottom panel.

Finally, we ran all three populations through the OOA3 model, which was originally described in [40] and recently implemented in `stdpopsim` [32]. This required using a 3-population CNN discriminator, which contains many more weights to optimize relative to the two population CNN. In addition, the OOA3 model requires 14 parameters. We inferred 10 of these parameters, fixing the four migration rate parameters and running `pg-gan` for 500 iterations. We also changed the mutation rate from 2.35 *×* 10^*−*8^ (which was used in [40]) to 1.29 *×* 10^*−*8^ (the recommended human mutation rate from [32]). The inferred parameters and summary statistics are shown in Figure S8, as well as a diagram of the demographic model (reproduced from Figure 2B in [40]). We find a discriminator accuracy of 0.65 and fit some but not all statistics well, suggesting model misspecification, or the difficulty of exploring a relatively high dimensional parameter space.

### Computational Resources

The runtime of our method is around 5-6 hours using a Quadro P5000 GPU. Pre-processing the real data takes several hours for each set of populations (YRI, CEU, CHB, YRI/CEU, YRI/CHB, CEU/CHB, and YRI/CEU/CHB). The resulting file sizes are 540M-944M, but these do not need to be loaded into memory due to the HDF5 format. The runtime for `fastsimcoal` was around 55 minutes.

## Discussion

We present a method for automatically learning parameters that can be used to simulate realistic genetic data. Most existing methods optimize parameters to match summary statistics like the SFS. Our algorithm, `pg-gan`, is a more holistic approach, which finds parameters that generate data that are systematically indistinguishable from the input data, although in practice it also often matches the summary statistics.

Our generative adversarial framework simultaneously trains a generator to produce reasonable evolutionary parameters and a discriminator to distinguish real data from simulated. We use real data during training to make sure the simulations capture realistic genomic features. We demonstrate the use of our method in an isolation-with-migration simulation setting and create simulated data that mirrors three human populations individually, in pairs, and all together. The discriminator often achieves accuracy between 50% and 70%, indicating strong, albeit incomplete, confusion between the real and simulated data. The approach is highly flexible and can automatically fit any parameterized model to any genomic data. We anticipate it will be particularly useful for understudied populations or species, since any unknown parameters can be included in the model and learned.

Our approach yields a natural way of evaluating and refining simulation pipelines. If simulations are easily distinguished from real data, then the model is not producing realistic data. We easily reach essentially complete (50%) discriminator confusion and good summary statistic matching in simulations. But with real data, the fit is imperfect. This could be because there are features of the real data that our models do not include, for example false negatives and other genotyping errors, phasing errors, missing data, and inaccessible regions of the genome. Through changes to the generative model, it would be possible to incorporate these effects and evaluate their impact. To handle limited power to detect rare variants (likely why we see more singletons and rare variants in the simulations than the real data), we experimented with filtering a fraction of singletons from the simulations. This improved the results for YRI, but not for CEU or CHB. Such filtering could be more adaptive in a future iteration. Features such as missing data could be important in some contexts (see `ReLERNN` [11] for an example of how to handle missing data). In general such data quality related features are dangerous for our approach because they provide a way for the discriminator to easily distinguish real and simulated data. For example if the generative model had data missing at random but the real data are missing in a non-random fashion then the discriminator will use this signal for classification. It would be important to make the generative model sufficiently flexible that it could learn to replicate the distinguishing features of the real data.

Some subsets of populations were more difficult to fit than others. The CEU/CHB split proved particularly difficult for both `pg-gan` and `fastsimcoal`. Since data quality should be similar between populations, this probably indicates that our model does not include demographic features that are important for patterns of variation in these populations, for example archaic admixture or exponential growth. More generally our model ignores many important biological features, for example heterogeneity in the mutation rate and other parameters, and natural selection. We assumed that mutation and recombination rates were known, but they can easily be added as parameters to the generative model and inferred (Figure S9). Heterogeneity could be modeled by fitting a distribution from which to draw parameters, rather than a point estimate. Natural selection, which can bias estimates of demographic parameters [49], is more difficult to model. The effects of regions under strong positive selection or long-term balancing selection can be minimized by removing them from the training data. However, background selection affects the majority of the genome and completely restricting to “truly neutral” regions of the genome is impractical. One simple but somewhat unsatisfactory solution is to approximate the effects of background selection across the genome by scaling effective population sizes with a factor drawn from empirical estimates of the effect of background selection across the genome [50]. A better solution would be to estimate the distribution of selection coefficients as part of the model [51]. This requires a generator that can simulate selection, for example SLiM [21], but would be much more computationally intensive than the coalescent simulations in the current approach. Efficiently incorporating selection into the model is a key area for future development.

There are several areas of future exploration that involve algorithmic modifications. In our current implementation, the topology of the demographic model needs to be specified ahead of time. However, it would be possible to extend our method to explore a space of demographic models, which would allow both the topology and the model parameters to be learned automatically. Although we mitigate overfitting by selecting real data regions at random (as opposed to a fixed sliding window), it is still a concern for the discriminator due to the fundamental data imbalance. The amount of real data is fixed, but the number of simulated examples is unlimited. There are many ways to guard against overfitting neural networks, including regularization and architecture modifications. An important line of future research is to optimize the training procedure in the presence of limited real data.

Another asymmetry comes from the potentially different learning rates of the generator and discriminator. The training of both components needs to be balanced – if the discriminator learns the difference between real and simulated data too quickly, the generator might not have a chance to explore a parameter space that would actually cause confusion. On the other hand, if the discriminator learns too slowly, all generator updates might look equally confusing. It would be interesting to explore adaptively controlling the learning rate – slowing down either the generator or the discriminator as needed through fewer parameter proposals or mini-batches. Understanding the behaviour of the discriminator is itself an important area of future work, which could help us investigate alignment between its hidden layers and traditional summary statistics.

Some idea of the uncertainty in the parameter space can be obtained by looking at the distribution of replicate estimates. In principle this approach could be extended to provide bootstrap confidence intervals by fitting the model to resampled data. A more general approach would be to fix the discriminator and vary the generator parameters to identify the parameter space over which the discriminator has low accuracy.

Our approach could be incorporated into a transfer learning [52] framework. In transfer learning, the parameters of an ML model are initialized by training on a large dataset, then “fine-tuned” by training on a smaller number of examples from the target dataset. In our case, a large dataset like the 1000 Genomes could be used to find an initial guess for discriminator weights, then these weights could be fine-tuned using data with fewer regions or sequenced individuals. The evolutionary model could still be modified, as transfer learning would be used for the discriminator, not the generator. The only restriction would be that the number of populations would need to match between the larger dataset and the smaller dataset. The original learning on the larger dataset would primarily assist the discriminator in learning general features of genomic datasets – population-level specifics would be learned in the fine-tuning phase.

It is our hope that others will build upon this initial exploration into parametric GANs for population genetics. Future developments will include integrating more realistic features of real data, constructing bootstrap confidence intervals for parameter estimates, and applying our approach to non-human species. In terms of methodological development, we aim to integrate transfer learning and develop interpretative approaches for the CNN discriminator, in order to investigate alignment between its hidden layers and traditional summary statistics. Modern machine learning has proved to be powerful in many domains, and our work emphasizes that this is true for population genetics as well. However, machine learning in population genetics requires novel architectures, for example our parametric generator and multi-population CNN discriminator – innovations that will be useful for future development of ML methods in the field.

## Acknowledgments

The authors would like to thank Joe Cammisa for extensive computational support, and Ke Wang for assistance with comparing `pg-gan` to other methods. SM is funded in part by NIH grant R15HG011528 and IM is funded in part by NIH grant R35GM133708. MK was funded through a David Robbins ’83 Big Data/Social Change Lang Center Internship. The content is solely the responsibility of the authors and does not necessarily represent the official views of the National Institutes of Health or other funding sources.

