## Supplementary Material for "Automatic inference of demographic parameters using Generative Adversarial Networks"

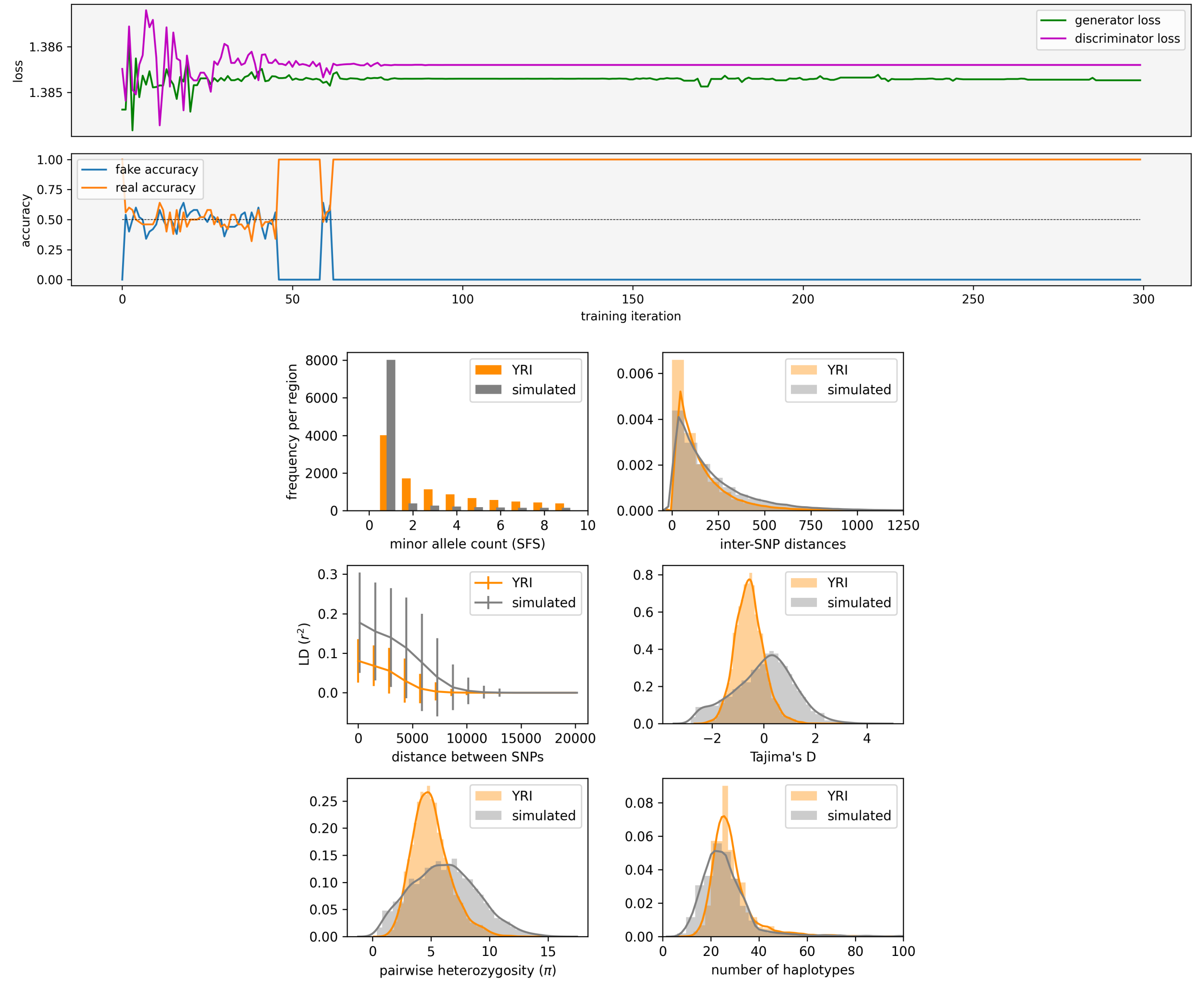


Figure S1: **YRI: failed run.** In this run the discriminator failed to learn and ended up predicting the same class for all regions (all real, so the real accuracy is 100% and the fake accuracy is 0%). This confuses the generator too, since regardless of the actual parameters, the generated data is classified as real. As a results, the inference becomes a random walk through the parameter space and the resulting summary statistics are far from the real data. **Top**: generator and discriminator losses across the training iterations (first panel), along with fake and real accuracy as output by the discriminator (second panel). **Bottom**: summary statistics under the resulting parameter inference, which are very far from the real data.

**IM model on simulated training data (with inferred mutation)**

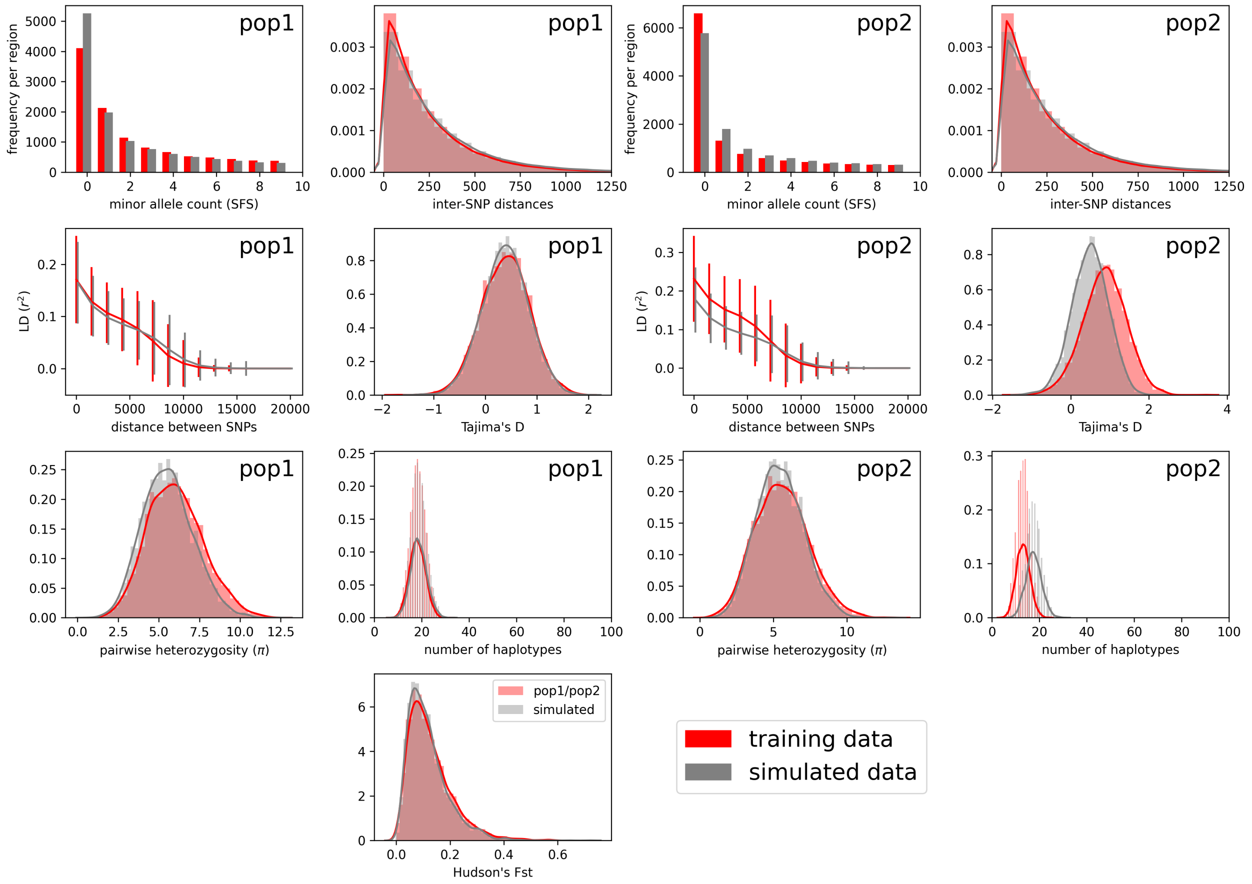


| | reco | mut | $N_{\text{anc}}$ | $T_{\text{split}}$ | mig | $N_1$ | $N_2$ |
|---|---|---|---|---|---|---|---|
| TRUE | 1.25e-08 | 1.25e-08 | 15000 | 2000 | 0.050 | 9000 | 5000 |
| `pg-gan` | 1.40e-08 | 8.77e-09 | 17568 | 2785 | -0.025 | 10580 | 9602 |

Figure S2: **IM model statistics on simulated training data (with mutation).** Summary statistics for data simulated under our inferred parameters ("simulated data"), compared with data simulated under the true parameters ("training data"). Subfigures on the left correspond to statistics from the first population, and those on the right correspond to the second population. In the bottom panel we show $F_{\text{st}}$ between the two populations. In this analysis we also infer the mutation rate (mut), along with the original six IM model parameters (see table above). Our results are close to the true parameters, except in the case of migration rate (mig) and $N_2$ – these discrepancies are reflected in the summary statistics for the second population.

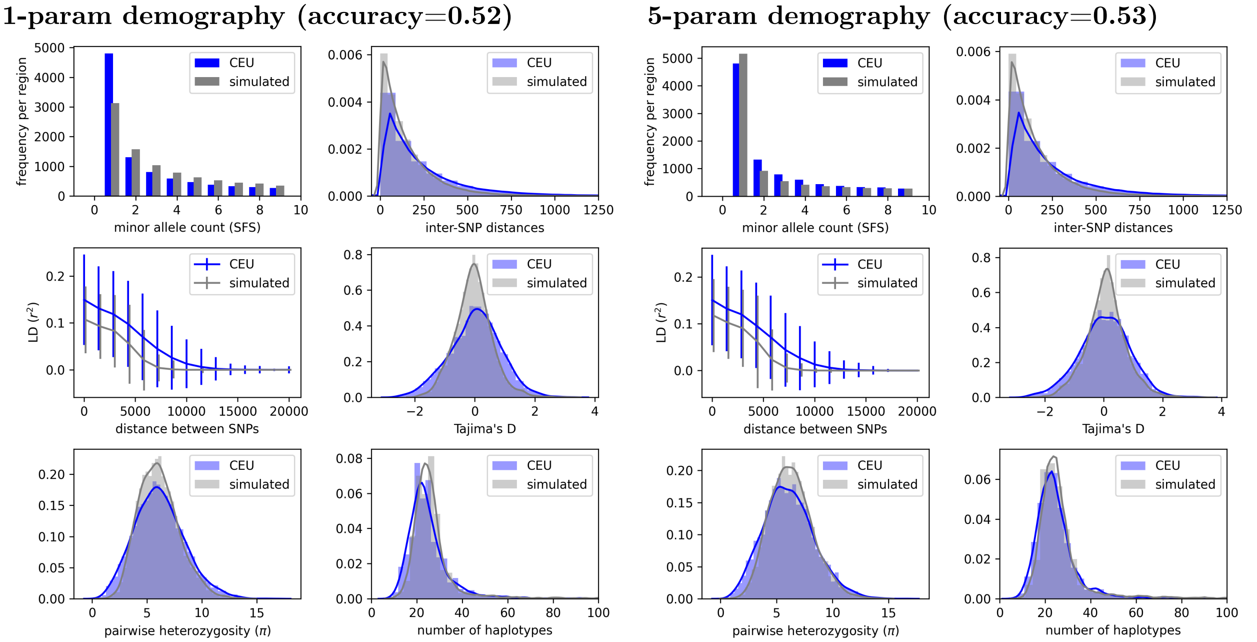


Figure S3: **CEU: one-population model.** Summary statistic comparison between CEU data from the 1000 Genomes project and simulated data under the inferred parameters from the two scenarios in the main text (see Figure 7). **Left**: simulated data under a constant population size with HapMap recombination rates. Simulated accuracy: 0.4, overall accuracy: 0.52. **Right**: Optimal model with `sum` permutation-invariant function, 5-parameter model, and HapMap recombination rates. Simulated accuracy: 0.6, overall accuracy: 0.53.

pg-gan: **YRI/CHB split (accuracy=0.63)**

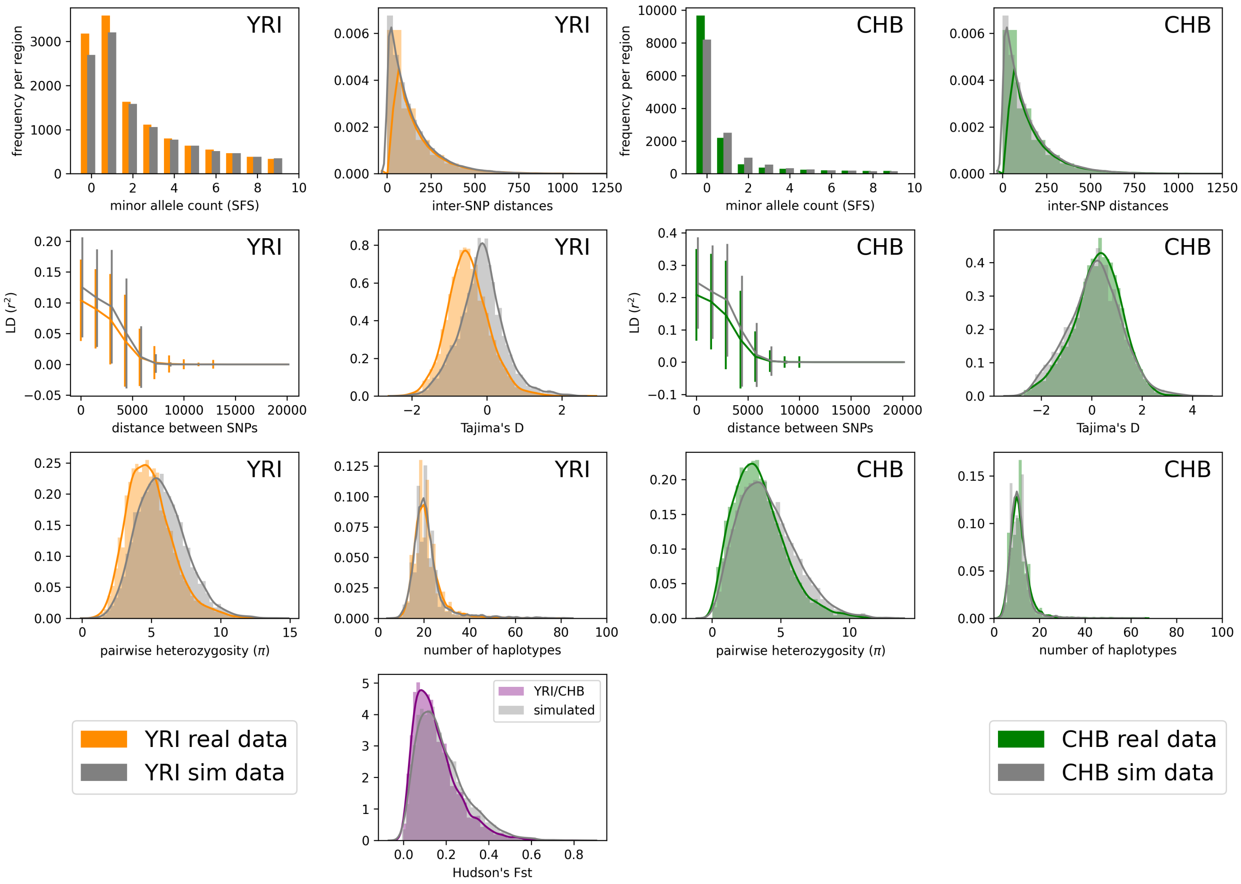


Figure S4: **YRI/CHB: two-population model (pg-gan).** Summary statistic comparison between real 1000 Genomes data and data simulated under the inferred parameters from pg-gan (see Table 3 for parameter values and Figure 3C for the OOA2 model). Left: statistics computed on YRI samples only. Right: statistics computed on CHB samples only. Note that we have non-segregating sites when considering each population separately, but not when we consider them together. $F_{\text{st}}$ between the two populations is shown in the last row. Simulated accuracy: 0.42, overall accuracy: 0.63.

**`fastsimcoal`: YRI/CHB split**

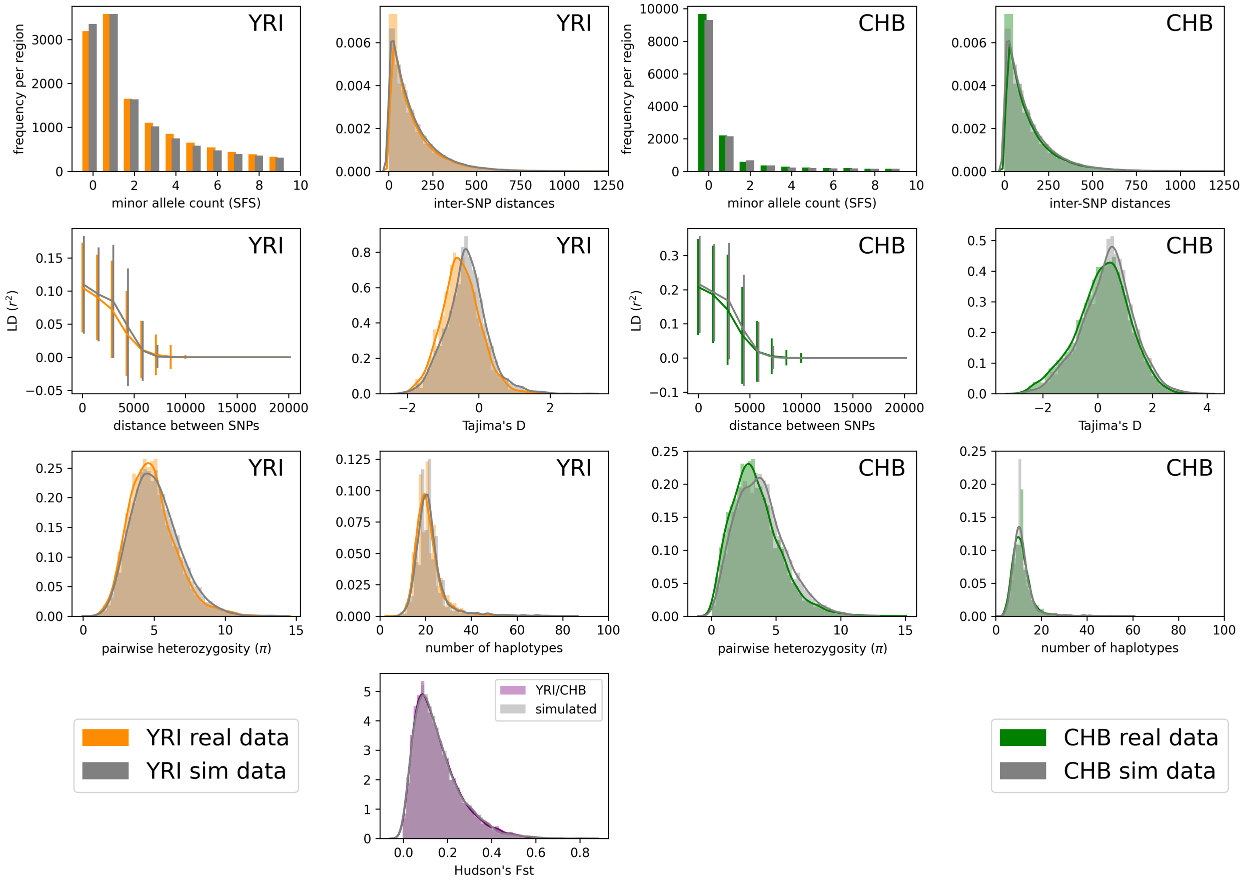


Figure S5: **YRI/CHB: two-population model (`fastsimcoal`).** Summary statistic comparison between real 1000 Genomes data and data simulated under the inferred parameters from `fastsimcoal` (see Table 3 for parameter values and Figure 3C for the OOA2 model). Left: statistics computed on YRI samples only. Right: statistics computed on CHB samples only. Note that we have non-segregating sites when considering each population separately, but not when we consider them together. $F_{\mathrm{st}}$ between the two populations is shown in the last row.

**pg-gan: CEU/CHB split (accuracy=0.77)**

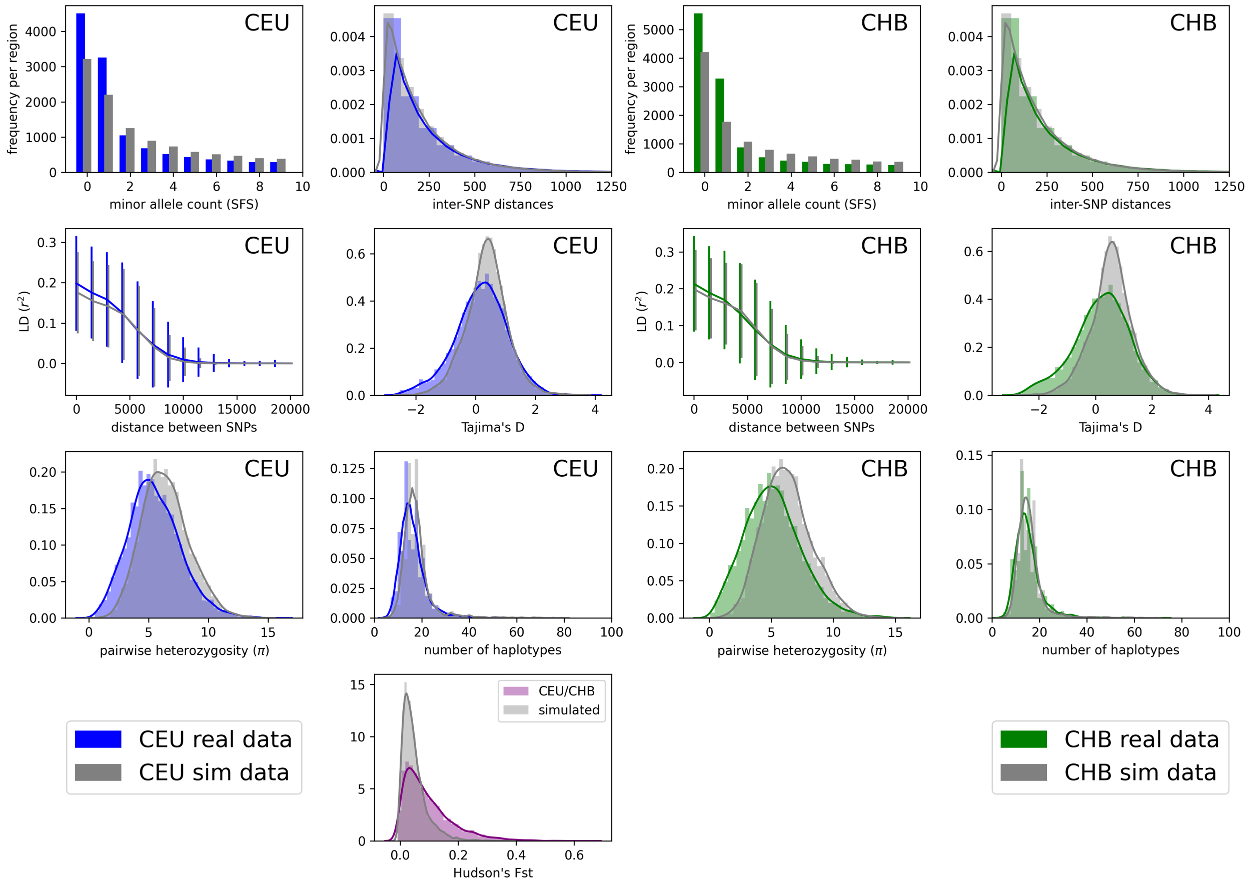


Figure S6: **CEU/CHB: two-population model.** Summary statistic comparison between real 1000 Genomes data and data simulated under the inferred parameters from `pg-gan` (see Table 3 for parameter values and Figure 3D for the POST model). Left: statistics computed on CEU samples only. Right: statistics computed on CHB samples only. Note that we have non-segregating sites when considering each population separately, but not when we consider them together. $F_{\mathrm{st}}$ between the two populations is shown in the last row, which we note is much less closely matched than for YRI/CEU or YRI/CHB. Simulated accuracy: 0.78, overall accuracy: 0.77.

**`fastsimcoal`: CEU/CHB split**

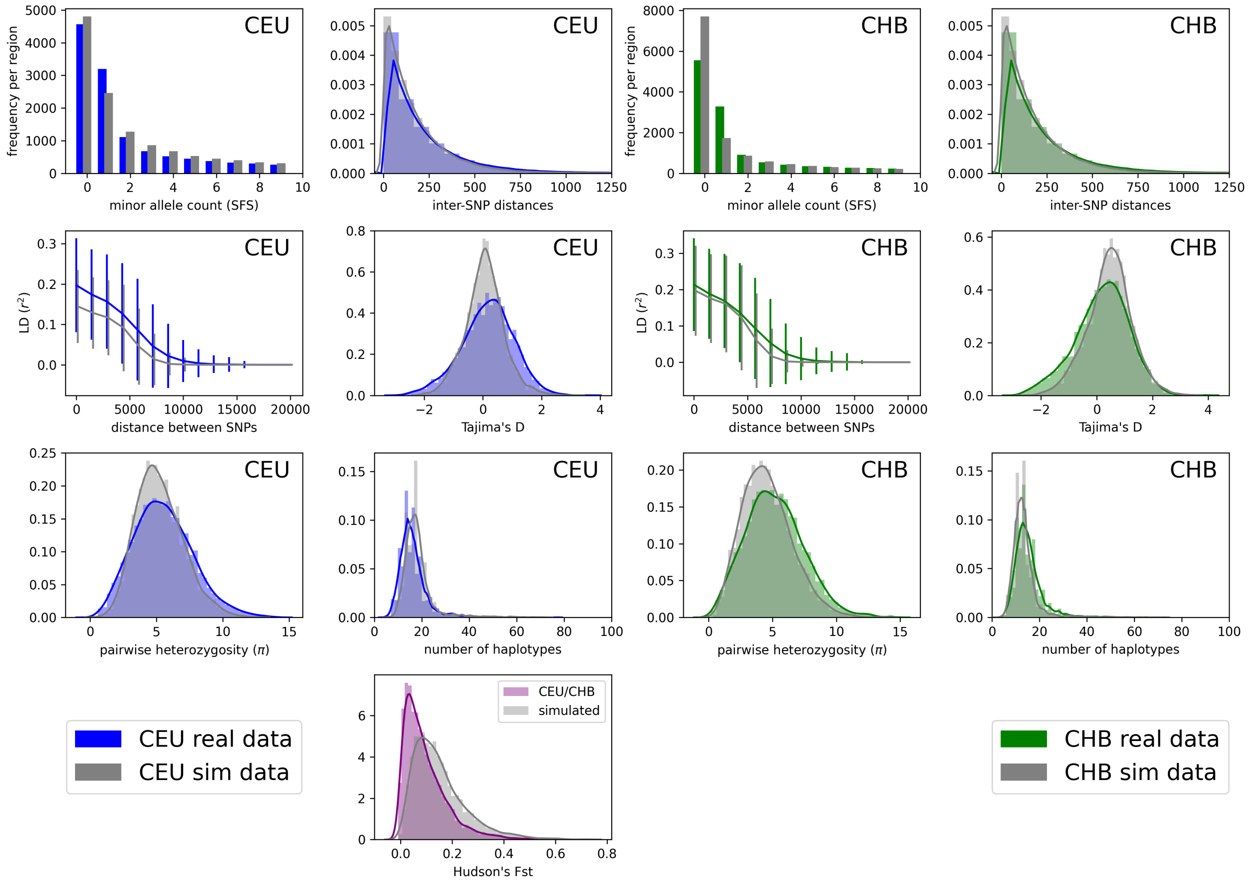


Figure S7: **CEU/CHB: two-population model (`fastsimcoal`).** Summary statistic comparison between real 1000 Genomes data and data simulated under the inferred parameters from `fastsimcoal` (see Table 3 for parameter values and Figure 3D for the POST model). Left: statistics computed on CEU samples only. Right: statistics computed on CHB samples only. Note that we have non-segregating sites when considering each population separately, but not when we consider them together. $F_{\text{st}}$ between the two populations is shown in the last row.

**pg-gan: YRI/CEU/CHB from OOA3 (accuracy=0.65)**

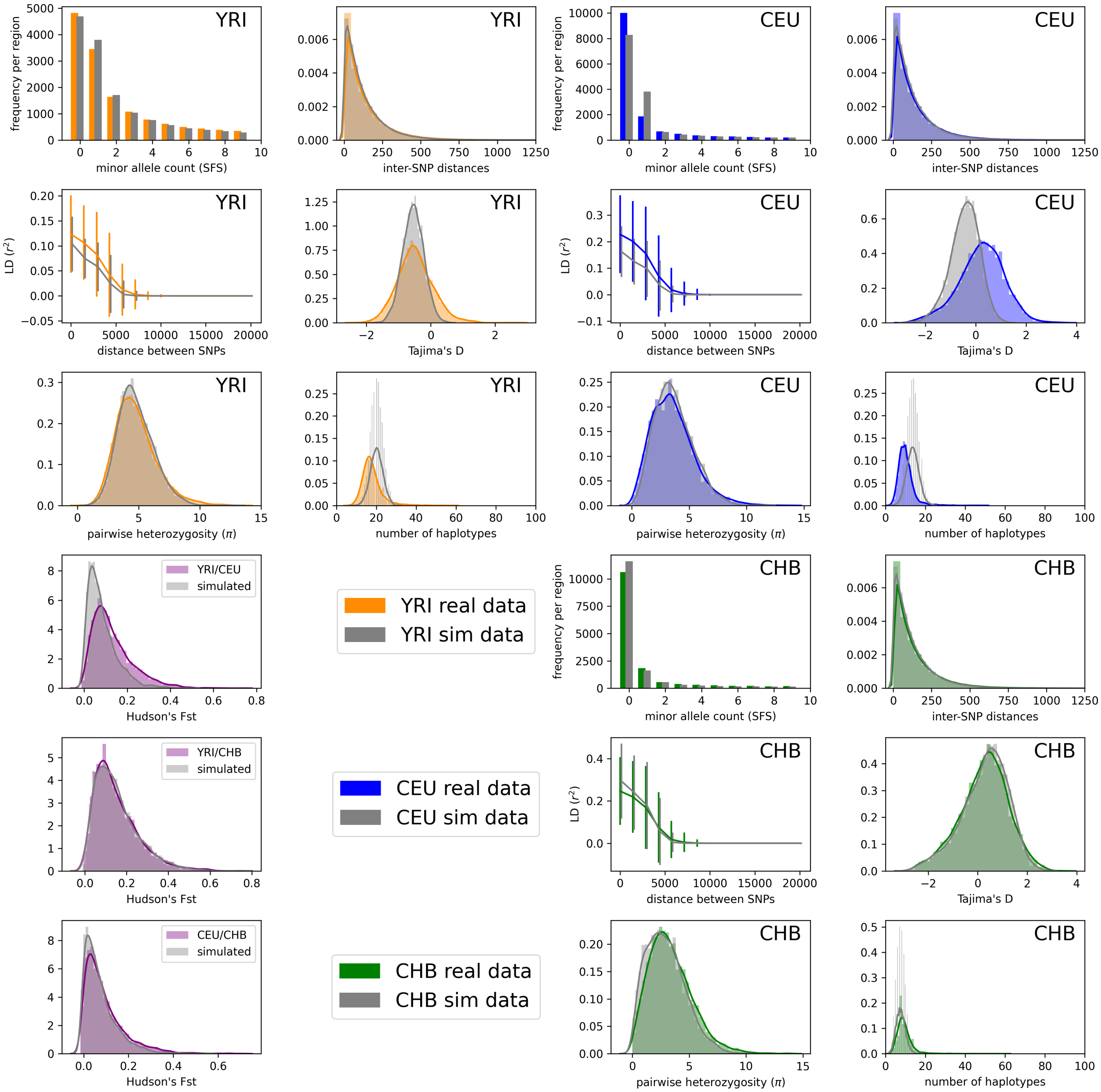


| Parameter | min | max | units | pg-gan inference |
|---|---|---|---|---|
| $N_A$ | 1000 | 30000 | individuals | 19,227 |
| $N_B$ | 1000 | 20000 | individuals | 2,174 |
| $N_{AF}$ | 1000 | 40000 | individuals | 36,247 |
| $N_{EU0}$ | 100 | 20000 | individuals | 16,723 |
| $N_{AS0}$ | 100 | 20000 | individuals | 576 |
| $r_{EU}$ | 0.0 | 0.05 | fraction of individuals | 0.0260 |
| $r_{AS}$ | 0.0 | 0.05 | fraction of individuals | 0.0030 |
| $T_{AF}$ | 8000 | 15000 | generations | 12,111 |
| $T_B$ | 2000 | 8000 | generations | 4,925 |
| $T_{EU-AS}$ | 100 | 2000 | generations | 1,325 |

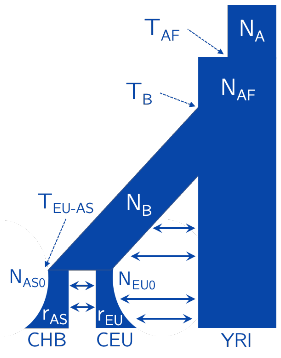


Figure S8: **YRI/CEU/CHB: three-population model.** Summary statistic comparison between real 1000 Genomes data and data simulated under the inferred parameters from the YRI/CEU/CHB OOA3 model with 10/14 parameters inferred and migration parameters fixed. The table below includes the ranges and inferred values for these 10 parameters. See [1] for the original specification of the OOA3 model and [2] for an implementation of the model in `msprime` [3]. Simulated accuracy: 0.68, overall accuracy: 0.65.

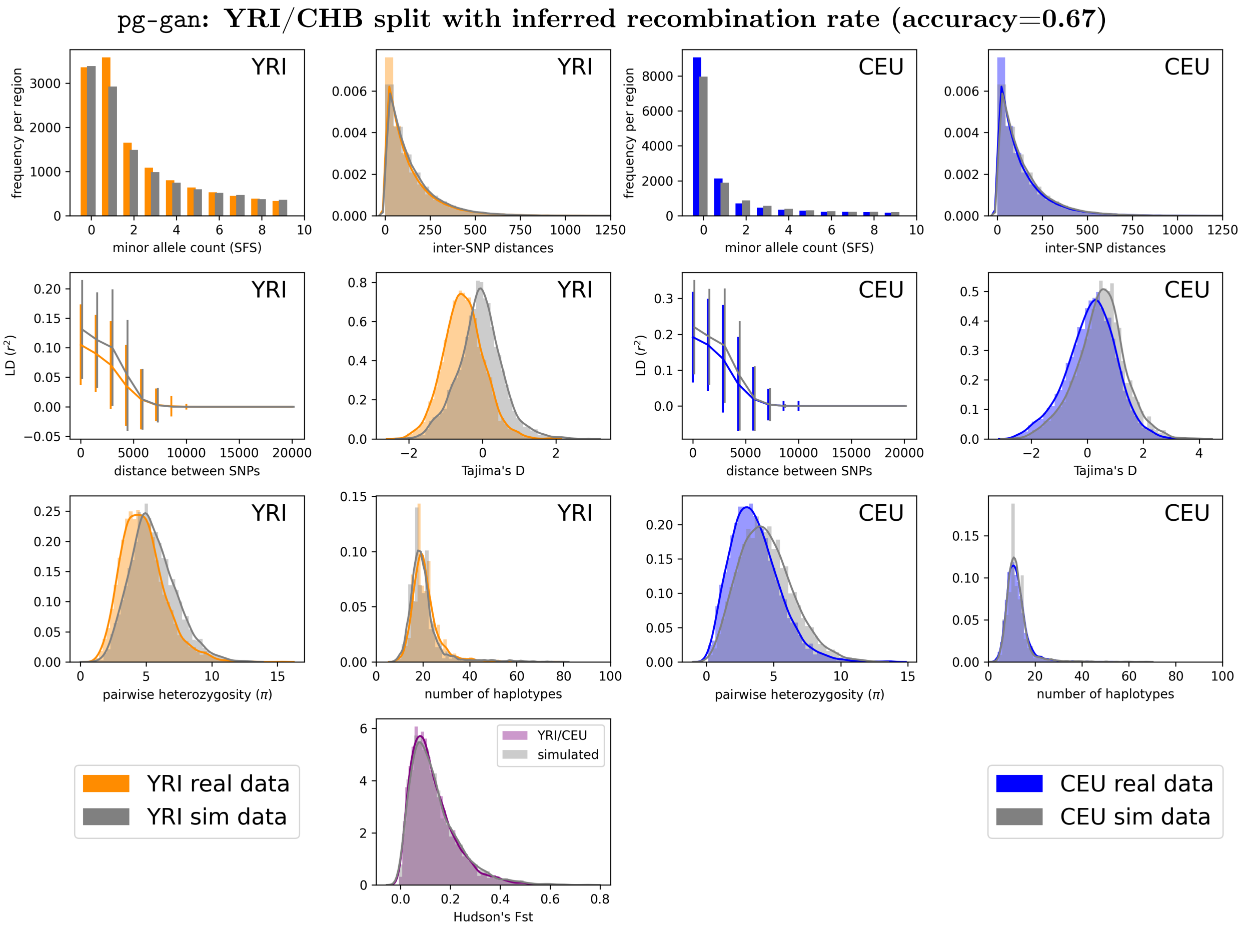


Figure S9: **YRI/CEU split with recombination rate inferred.** In this analysis we infer the recombination rate instead of sampling from the HapMap recombination rate. We inferred a rate of 1.20e-08 in this scenario. The fit of the summary statistics is not as close as when we use the HapMap rates (see Figure 9). Simulated accuracy: 0.52, overall accuracy: 0.67.
